# Discovering Prognostic Risk Strata and Heterogeneous Progression Routes from Cross-Sectional Survival Data

**DOI:** 10.64898/2026.11.02.763853

**Authors:** Ethar Alzaid, George Wright, Fayyaz Minhas

**Author notes:** Contributing authors.

## Abstract

Inferring how individual patients progress towards higher disease risk typically requires longitudinal observations, which are often unavailable. Population-level analyses of cross-sectional data can identify differences between risk groups but may obscure heterogeneous progression patterns across individual patients. In this work, we introduce GOTSurv, a framework for patient-specific progression modelling. Using cross-sectional survival data, it first learns to rank patients by survival risk and automatically stratify them into prognostic risk groups without requiring predefined risk cut-offs. It then uses unbalanced optimal transport to infer patient-specific changes in the magnitude and direction of features associated with transitions towards higher-risk strata. We evaluated GOTSurv across ten TCGA cancer cohorts using two modalities: gene expression (n=4,767) and whole-slide image representations (n=4,655), with external validation in two independent cohorts. Compared with established survival models, GOTSurv generally achieved stronger prognostic stratification without requiring predefined risk cut-offs, while retaining competitive survival-ranking accuracy. More importantly, GOTSurv enabled the discovery of individualised progression trajectories associated with increasing risk, revealing distinct progression patterns even among patients within the same risk stratum. Overall, GOTSurv provides a novel framework for inferring heterogeneous, patient-specific progression patterns from cross-sectional survival data, enabling the identification of distinct pathways associated with increasing disease risk and informing the development of personalised therapeutic strategies. The code for GOTSurv is available at https://github.com/EtharZaid/GOTSurv/.

## 1 Introduction

Disease progression modelling aims to understand how patients move from one disease state or risk level to another, and which changes in their observed data are associated with progression towards higher risk [1]. Understanding how disease patterns progresses in an individual patient is important for personalised patient care and informing treatment planning and patient monitoring [2–4]. However, accurately modelling progression at the individual-patient level typically requires longitudinal data, which are often unavailable.

A common approach to understanding the changes associated with disease progression from cross-sectional data is to first use a risk prediction model to stratify patients into two risk groups (low v. high) [5–7], and then compare the expression profiles of these groups using conventional analyses such as differential expression. However, such analyses provide only a population-level summary and do not capture how an individual patient’s profile may change as they progress towards higher risk.

Population-level summaries do not necessarily reflect individual behaviour, particularly in heterogeneous populations, such as cancer cohorts where patients within the same risk group may differ substantially in their molecular profiles and disease pathways. Studies of clinical outcomes and gene-expression data have shown that group-level effects can mask substantial inter-individual variation, highlighting the need for analyses at the individual-patient level [8–11].

One way to capture this heterogeneity is to model progression at the level of individual samples rather than only comparing populations. Methods for inferring progression from unpaired samples have been widely explored, particularly in singlecell biology, where approaches reconstruct transitions between cellular states from cross-sectional observations [12–14]. Related work has also considered patient-level progression from cross-sectional data [15–17]. While these methods can model progression for individual patients based on disease stage or subtype, to the best of our knowledge, no existing method jointly discovers prognostic risk groups and models how individual patients’ features evolve as they progress towards higher-risk disease states.

In this work, we introduce GOTSurv, a framework that uses cross-sectional patient data to automatically discover risk groups and infer risk-specific progression trajectories for individual patients. Each trajectory describes how a patient’s features are inferred to change from their current risk level towards the next higher-risk group, where risk levels are defined by survival outcomes. Although progression is modelled between risk groups, the resulting trajectory is estimated for each patient individually. Patients within the same risk level can therefore follow different trajectories associated with distinct molecular profiles and pathways, preserving inter-individual variation that may be obscured by population-level comparisons. To our knowledge, GOTSurv is the first method to model how an individual patient’s features change towards higher survival risk from cross-sectional data.

The main contributions of this work are as follows:

1. We introduce GOTSurv, a framework that automatically discovers prognostic risk strata from cross-sectional survival data and uses them to infer patient-specific progression trajectories towards higher-risk states, enabling the discovery of heterogeneous molecular progression patterns without requiring longitudinal observations.
2. We demonstrate that GOTSurv reveals heterogeneity within the same risk level by identifying subgroups of patients with distinct inferred progression patterns, characterised by differences in molecular profiles and pathway activity that are not explained by risk alone.
3. We benchmark GOTSurv against established survival models across 10 TCGA cancer cohorts using gene-expression and whole-slide image representations, demonstrating improved risk stratification, discovery of heterogeneous progression trajectories, generalisability to independent external cohorts, and uncertainty-aware predictions through selective abstention.

## 2 Methods

### 2.1 Problem Formulation and Overview

We consider a cross-sectional dataset D comprising patients represented by *d*-dimensional feature vectors **x**_**i**_ ∈ ℝ^*d*^, which may encode molecular measurements, imaging-derived characteristics, or any other fixed-length representation. We partition the dataset into a labelled subset ℒ, containing patients with known survival outcomes (*t*_*i*_, *e*_*i*_), and an unlabelled subset *U*, for which survival outcomes are unavailable, such that D = ℒ ∪ *U*. Here, *t*_*i*_ ∈ ℝ+ denotes the observed follow-up time and *e*_*i*_ ∈ 0, 1 indicates whether the event of interest was observed (*e*_*i*_ = 1) or the observation was censored (*e*_*i*_ = 0).

Our aim is to characterise how individual patients may progress through the prognostic risk space by identifying the patient-specific changes associated with transitions from lower to higher risk. To achieve this, GOTSurv performs two complementary tasks, as illustrated in Fig. 1A. First, unlike conventional survival analysis and risk-stratification approaches, which often rely on predefined risk thresholds, percentile-based cut-offs, or post hoc grouping of predicted risk scores, GOTSurv jointly learns survival-risk rankings and discovers *K* prognostic groups with progressively increasing risk in a fully data-driven manner. Second, rather than limiting the analysis to risk prediction or group assignment, GOTSurv learns mappings between these ordered risk groups to infer patient-specific progression vectors using optimal transport. These vectors characterise the changes associated with movement towards higher-risk states, enabling downstream analyses aimed at identifying the biological features underlying individualised risk progression. We describe these two steps in detail below.

**Fig. 1.**
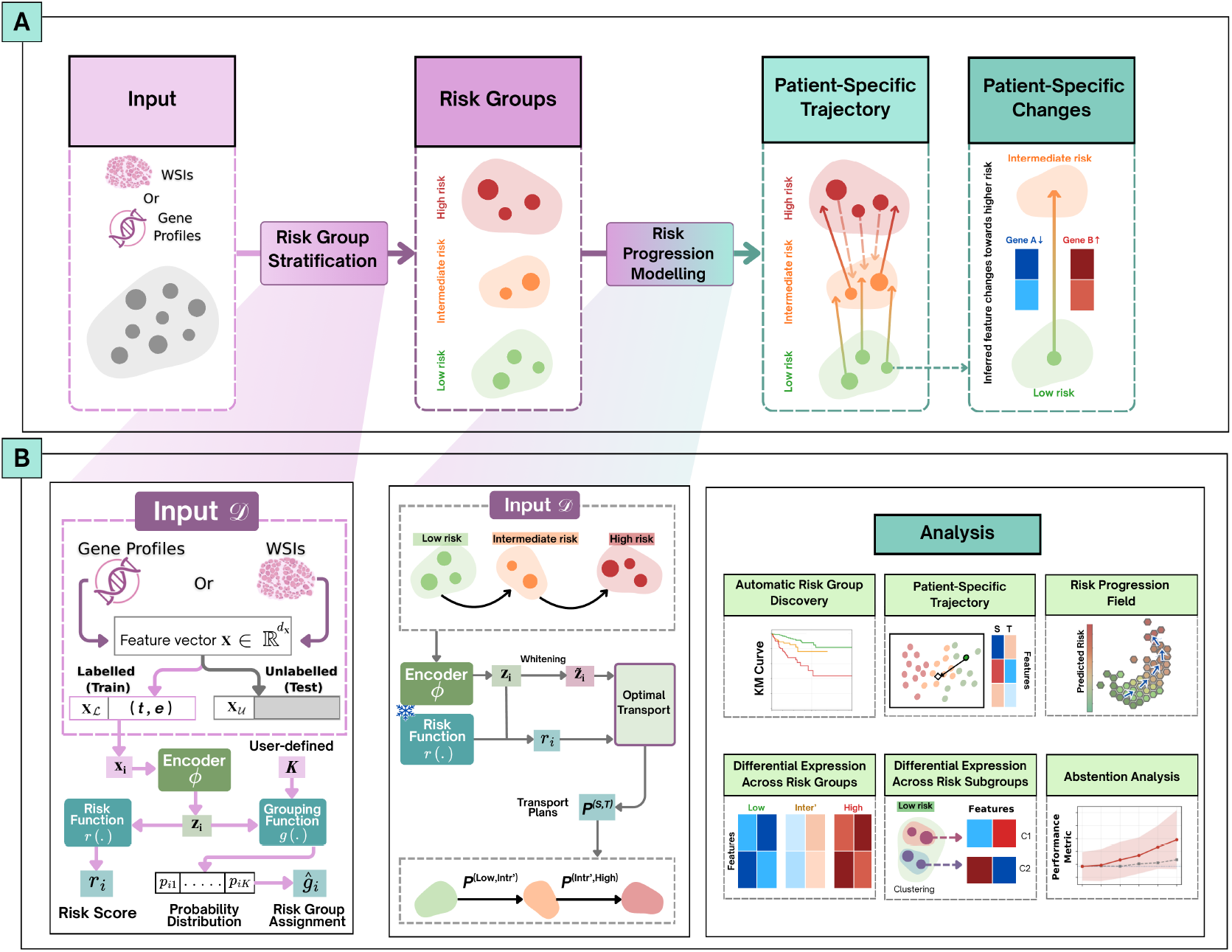
Overview of GOTSurv. **(A)** Conceptual workflow. Cross-sectional patient data are stratified into ordered survival-risk groups, and progression is modelled between adjacent groups to derive patient-specific trajectories and the corresponding feature changes towards higher risk. **(B)** Detailed framework. Labelled samples are used to jointly learn a latent representation, survival-risk score, and risk-group assignments, which are then applied to the full dataset. The learnt representation is carried forward to the progression-modelling stage, where optimal transport estimates transport plans between adjacent risk groups. These outputs support downstream analyses of risk stratification, patient-specific trajectories, progression fields, and molecular differences both across and within risk groups.

### 2.2 Automatic Risk-Based Patient Stratification and Confidence Estimation

Our first goal is to stratify patients into *K* prognostic groups with progressively increasing risk in a fully data-driven manner, without requiring predefined risk thresholds or percentile-based cut-offs, such that patients assigned to higher-indexed groups are expected to experience adverse events earlier than those in lower-indexed groups. To this end, GOTSurv jointly learns a continuous survival-risk score and a probability of belonging to each of the *K* risk groups for every patient, by minimising the expected ranking error over comparable patient pairs. The resulting membership probabilities also provide a measure of assignment confidence, enabling the model to abstain from uncertain predictions.

Each patient *i*, represented by a feature vector **x**_*i*_, is mapped to a latent representation **z**_*i*_ = *ϕ*(**x**_*i*_; *θ*_*ϕ*_), from which we predict a continuous risk score *r*_*i*_ = *r*(**z**_*i*_; *θ*_*r*_) and the probability *p*_*ik*_ = *g*_*k*_(**z**_*i*_; *θ*_*g*_) of belonging to risk group *k* ∈ {1, …, *K*}, where *p*_*ik*_ ∈ [0, 1] and 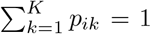. The functions *ϕ, r*, and *g* are neural networks heads with learnable parameters *θ*_*ϕ*_, *θ*_*r*_, and *θ*_*g*_, respectively. These parameters are jointly optimised using comparable patient pairs for which a relative risk ordering can be inferred from the observed survival times and event indicators. The set of such pairs is defined as

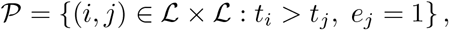

where patient *j* experienced an event while patient *i* remained event-free for a longer period, indicating that patient *j* is expected to have a higher risk than patient *i*. To quantify the discrepancy between these survival-derived risk orderings and the predicted risk scores, we define an ordering loss as the expected pairwise ranking error under a probability distribution *π* over comparable pairs:

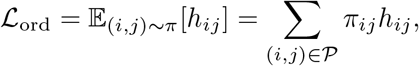

where *h*_*ij*_ = [1 − (*r*_*j*_ − *r*_*i*_)]_+_ is a margin-based ranking penalty, with [*x*]_+_ = max(0, *x*). This penalty measures whether the predicted risk difference between *j* and *i* satisfies the required survival-ordering margin. The probability distribution *π* is determined by how consistently the predicted risk-group memberships reflect the observed survival ordering. Specifically, for each comparable pair (*i, j*) ∈ P, we first define the groupordering agreement probability *a*_*ij*_

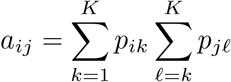

which represents the probability that the earlier-event patient *j* belongs to the same or a higher-indexed risk group than patient *i*. These agreement probabilities then define a distribution over comparable pairs through normalisation:

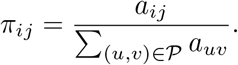

In the ordering loss, *π*_*ij*_ represents the probability of selecting pair (*i, j*) and determines its the weight of the overall ranking error over these pairs. This creates a bidirectional coupling in the minimisation of ordering loss: while group memberships influence risk-score learning through *π*_*ij*_, the resulting ranking errors also guide the learning of group memberships through *a*_*ij*_. This interaction between group discovery and risk score assignment is characterised by two complementary gradients (see Supplementary Methods for their derivations):

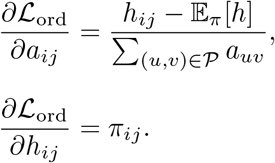

The first gradient describes how ranking performance influences group discovery. Conceptually, a gradient descent update 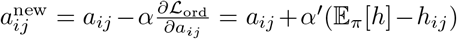, where *α*^*′*^ *>* 0 is the learning rate, favours increasing *a*_*ij*_ when the pairwise ranking error *h*_*ij*_ falls below the expected ranking error E_*π*_[*h*] across all comparable patient pairs under the distribution *π*. This encourages group memberships in which the earlierevent patient *j* is more likely to belong to the same or a higher-indexed risk group than patient *i*.

Conversely, the second gradient shows how group memberships influence risk prediction. For a pair with an active ranking penalty (*h*_*ij*_ *>* 0), a conceptual gradient descent update of the risk scores gives 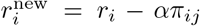 and 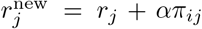. Thus, higher group-ordering agreement *a*_*ij*_ results in a stronger gradient contribution that increases the predicted risk of the earlier-event patient *j* relative to patient *i*, thereby reducing the pairwise ranking error and promoting stratification. These gradients show how survival-risk prediction and group discovery reinforce each other during optimisation, enabling risk groups to be learnt without predefined thresholds or boundaries.

To discourage group collapse, we introduce a minimum group-proportion penalty,

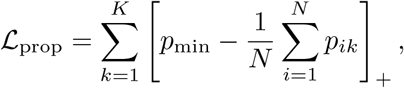

where *p*_min_ specifies the minimum desired proportion of patients assigned to each group, measured through their probabilistic memberships, and *N* is the number of patients used to evaluate the penalty. This penalises groups whose expected share of patients falls below *p*_min_, discouraging concentration of membership probability in only a few groups.

We additionally apply *L*_2_ regularisation to the weight parameters of the riskprediction network, defined as 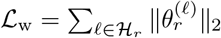, where H denotes the set of linear layers and 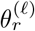 represents the weight matrix of layer *l*. The complete stratification objective is

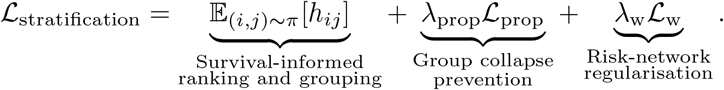

where *λ*_prop_ controls the contribution of the minimum group-proportion penalty and *λ*_w_ controls the strength of the regularisation. We fixed *λ*_prop_ = 0.65 and *λ*_w_ = 0.1 across all cohorts and modalities. These values were selected in preliminary experiments as a practical trade-off between preventing group collapse and preserving survival-ranking performance, and were not tuned separately for individual datasets.

Following optimisation, each patient is assigned to the highest-probability group, ĝ_*i*_ = arg max_*k*_ *p*_*ik*_, and groups are ordered by mean predicted risk so that higher indices indicate higher risk, ensuring that higher group indices correspond to higher average predicted risk. Figure 1B summarises how these assignments are carried forward into progression modelling. We also define assignment confidence as *c*_*i*_ = max_*k*_ *p*_*ik*_, with higher values indicating more decisive group membership. This score is used for selective abstention of low-confidence patients, thereby improving the reliability of risk stratification for the retained patients.

### 2.3 Progression Modelling

GOTSurv next uses the predicted risk-group assignments of all patients in D, including both labelled and unlabelled samples, to infer how the population at risk level *k* may transition towards the next higher-risk population *k* + 1. To enable this over cross-sectional data, we formulate this using Optimal transport (OT) [18], which models how one distribution can be transformed into another by finding the most efficient way to move mass between them. In our setting, this step produces a transport plan for each consecutive pair of risk groups (*k* → *k* + 1), describing how patients in the lower-risk population are mapped to patients in the higher-risk population based on their learnt representations. These transport plans provide the basis for inferring patient-specific progression vectors from cross-sectional data. We describe the technical details of this process below.

We organise the risk groups into a linear progression graph *G*, with directed edges connecting consecutive groups from lowest to highest risk. Each edge is denoted by (*S, T*), where *S* and *T* represent the source and target risk groups, respectively. For example, when *K* = 3, the graph comprises a transition from low to intermediate risk (*k* = 1 → *k* = 2) and another from intermediate to high risk (*k* = 2 → *k* = 3).

The first step in learning the transport mappings is to obtain a suitable latent representation of the input features. To this end, we reuse the encoder *ϕ* learnt during risk stratification, initialising it with the previously learnt parameters *θ*_*ϕ*_ to obtain latent representations **z**_*i*_ = *ϕ*(**x**_*i*_; *θ*_*ϕ*_), which are further refined during progression modelling. However, directly minimising transport costs can cause these representations to collapse towards similar values. To mitigate this, we apply covariance whitening: 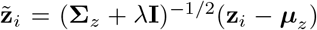, where ***µ***_*z*_ and **Σ**_*z*_ are the mean and covariance of the embeddings and *λ >* 0 is a small constant for numerical stability.

To formulate transport between risk groups, we represent each source and target group as an empirical probability distribution over the whitened latent representations. For each edge (*S, T*), let *S* and *T* denote the sets of patients assigned to the source and target groups, respectively. Assuming equal importance for all patients within each group, we assign uniform probability masses *a*_*i*_ = 1*/*|*S*| and *b*_*j*_ = 1*/*|*T* |. The resulting empirical distributions are defined as

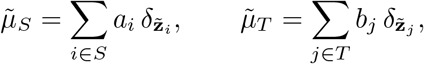

where 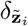 denotes a Dirac measure centred at 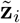.

The transport between the two a source and target set is represented by a transport plan 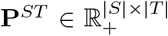, where each entry 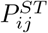 denotes the amount of probability mass transported from patient *i* ∈ *S* to patient *j* ∈ *T*. For each edge, we define the unbalanced OT objective

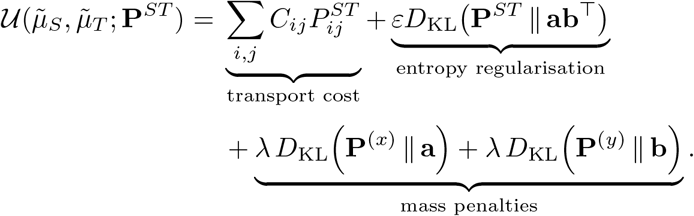

Here, 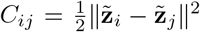 is the transport cost between patients in the learnt feature space, while 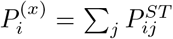 and 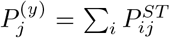 denote the transported source and target masses, respectively, and *D*_KL_ is the generalised Kullback–Leibler divergence. The transport-cost term favours matching patients with similar representations. The parameter *ε* controls how widely transport mass is distributed across target patients. Larger values spread mass across more patients in the next higher-risk stratum, while smaller values favour fewer, lower-cost correspondences. The parameter *λ* controls the strength of mass preservation between risk strata. Smaller values allow patients with high transport costs to remain effectively unmatched, while larger values encourage matching even when transport costs are high.

GOTSurv jointly learns the shared latent representation and the edge-specific transport plans across all edges in the progression graph through a nested optimisation:

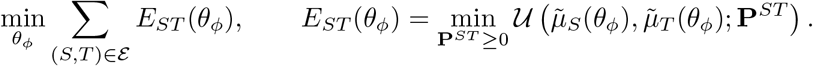

Here, ℰ denotes the set of directed edges in *G*, and *E*_*ST*_ (*θ*_*ϕ*_) is the OT energy obtained by minimising the previously defined unbalanced OT objective *U*. For the current encoder parameters, the inner optimisation estimates the transport plans using generalised Sinkhorn iterations, with the standard debiasing correction applied to the resulting transport energies to remove self-interaction bias introduced by entropy regularisation [19]. The outer optimisation then updates the shared encoder parameters through gradient descent using the debiased transport energies across all edges. As the encoder changes, the transport costs and corresponding plans are updated, allowing the latent representation and transport mappings to be learnt jointly.

The transport plans couple patients to the next higher-risk group and are used to derive patient-specific progression vectors. For patients in the highest-risk group, which has no outgoing transition, the final incoming transport plan is instead used to characterise their correspondences with patients in the preceding risk group. The complete GOTSurv training procedure, from risk-group discovery to optimisation of the transport mappings, is summarised in Algorithm 1.

In the implementation, *ε* and *λ* are parameterised as 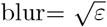 and 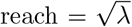, respectively. We fixed blur = 0.5 and reach = 0.01 across all cohorts and modalities, without dataset-specific tuning. A sensitivity analysis varying blur and reach around these values is provided in the Supplementary, confirming that inferred progression are stable across a broad range of both parameters.

### 2.4 Sample-Specific Progression Vectors

Beyond group-level transitions, the transport plans provide patient-to-patient correspondences across adjacent risk groups, enabling patient-specific progression vectors and associated feature changes to be inferred. For each source sample, we summarise its transport towards the target population to obtain a *sample-specific progression vector*. The objective of these vectors is to quantify how much of each feature has changed in order for a specific sample to move to higher-risk.

#### Algorithm 1

Overall GOTSurv training procedure

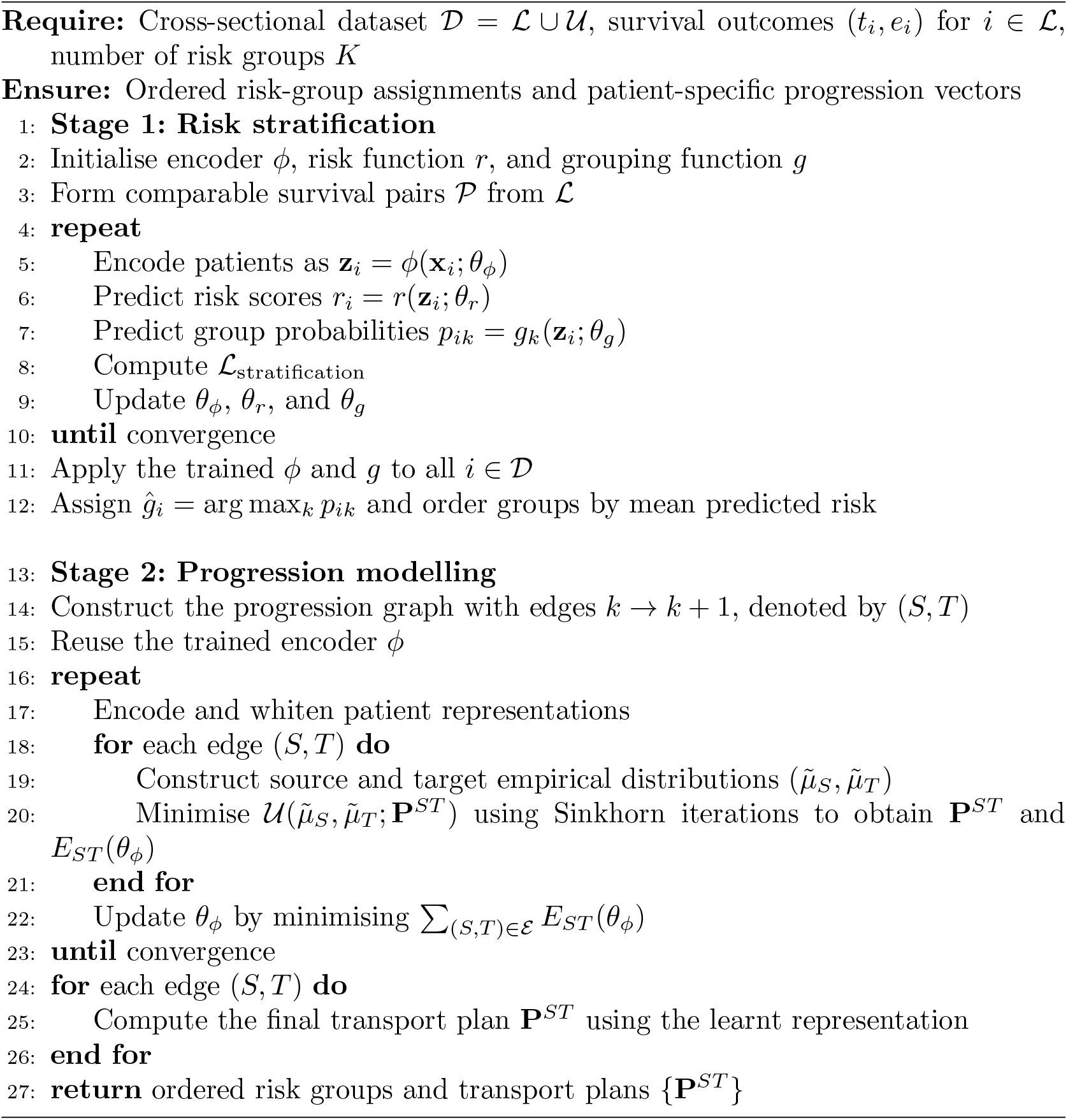

For a source sample *u* ∈ *S*, the transport plan does not generally pair it with a single target sample. Instead, its transported mass can be distributed across multiple samples in the target group. We therefore define a barycentric target as the transport-weighted average of the target samples, and the corresponding progression vector as the displacement from the source sample to this target,

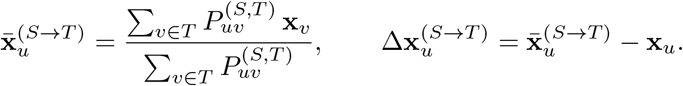

The barycentric target 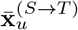 represents the point in the target population towards which source sample *u* is transported, while 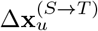 describes the inferred feature changes associated with its progression towards the target risk group.

For samples in the highest-risk group, this formulation cannot be applied directly because these terminal samples have no outgoing edge from which to infer further progression. Rather than discarding them, we use the transport plan from the final incoming edge, **P**^(*S,T*)^. For each terminal sample, we form a barycentric projection using the mass transported *into* that sample, allowing us to summarise the direction of change associated with its arrival in the terminal risk group.

Additionally, we derive latent-space progression vectors by replacing **x** with the whitened embedding 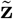. These are used for transport-field visualisation, while feature-space vectors support molecular interpretation, patient-target comparisons, and within-risk subgroup analysis.

### 2.5 Evaluation Protocol

We evaluated GOTSurv across ten TCGA cancer cohorts [20] using gene-expression and whole-slide image (WSI) representations, together with two independent external cohorts. We fixed the number of risk groups to *K* = 3 (low, intermediate, and high risk) across all experiments to provide a consistent progression structure across cohorts, sensitivity to *K* is evaluated in the Supplementary.

#### 2.5.1 Datasets

All modalities were represented as fixed-length feature vectors. For gene expression, cancer-specific prognostic gene sets were selected following Uhlén et al. [21], yielding 300–1,000 genes per cohort. WSIs were represented using 768-dimensional embeddings extracted with a pretrained slide-level model [22]. External validation used the CPTAC colorectal cancer cohort (*n* = 97) for gene expression [23] and the Australian Breast Cancer Tissue Bank (ABCTB; *n* = 2,311) for WSIs [24], using the same feature definitions as their corresponding TCGA cohorts.

#### 2.5.2 Evaluation Metrics

We evaluated survival ranking using the concordance index (C-index) [25]. Risk-group stratification was assessed using Kaplan–Meier (KM) curves and the multivariate log-rank test [26], together with restricted mean survival time (RMST) differences [27]. We report the adjacent-group RMST difference, corresponding to the neighbouring risk-group transitions modelled by GOTSurv.

#### 2.5.3 Baselines and Benchmarking Protocol

We benchmark GOTSurv against established survival models that operate on the same fixed-dimensional feature vectors. The models are: Cox proportional hazards regression (CoxPH) [28], gradient boosting machines (GBM) [29], random survival forests (RSF) [30], and survival support vector machines (SSVM) [31]. All models operated on the same input feature matrices and were evaluated over 100 bootstrap resamples using identical train–test splits. Baseline risk groups were obtained by tertile cut-points fitted to training-set risk scores and applied unchanged to the corresponding test samples, whereas GOTSurv used the groups produced directly by its grouping function. For external validation, models were trained on the full TCGA cohort and applied to the independent cohort without refitting or recalibration.

#### 2.5.4 Consensus Estimates

All analyses beyond the benchmark used out-of-sample predictions only. For each patient, predictions were aggregated across the resamples in which that patient appeared in the test set: mean predicted risk defined the consensus risk score, modal assignment defined the consensus risk group, and the mean progression vector defined the consensus progression vector.

## 3 Results

We organise the results around three main questions: how well GOTSurv predicts and separates risk, what progression it captures for individual patients, and whether patients within the same risk group follow different progression patterns.

### 3.1 Improved risk stratification with GOTSurv

Before using the learnt risk groups as states for progression modelling, we first assess whether they represent meaningful and ordered differences in survival. We therefore evaluated the automatically discovered low-, intermediate-, and high-risk groups using Kaplan–Meier survival curves (Fig. 2, KMs for the WSI cohorts are Supplementary Fig.S1). In CRC, the learnt risk groups showed clear and ordered survival separation. The RMST difference was 184 days between the low- and intermediate-risk groups and increased to 439 days between the low- and high-risk groups, with an intermediate-to-high difference of 255 days. This separation was highly significant (*p* = 1.3 × 10^−11^), indicating that the groups discovered directly from the data correspond to distinct survival outcomes. Importantly, this stratification was accompanied by strong ranking performance, with a C-index of 0.77.

**Fig. 2.**
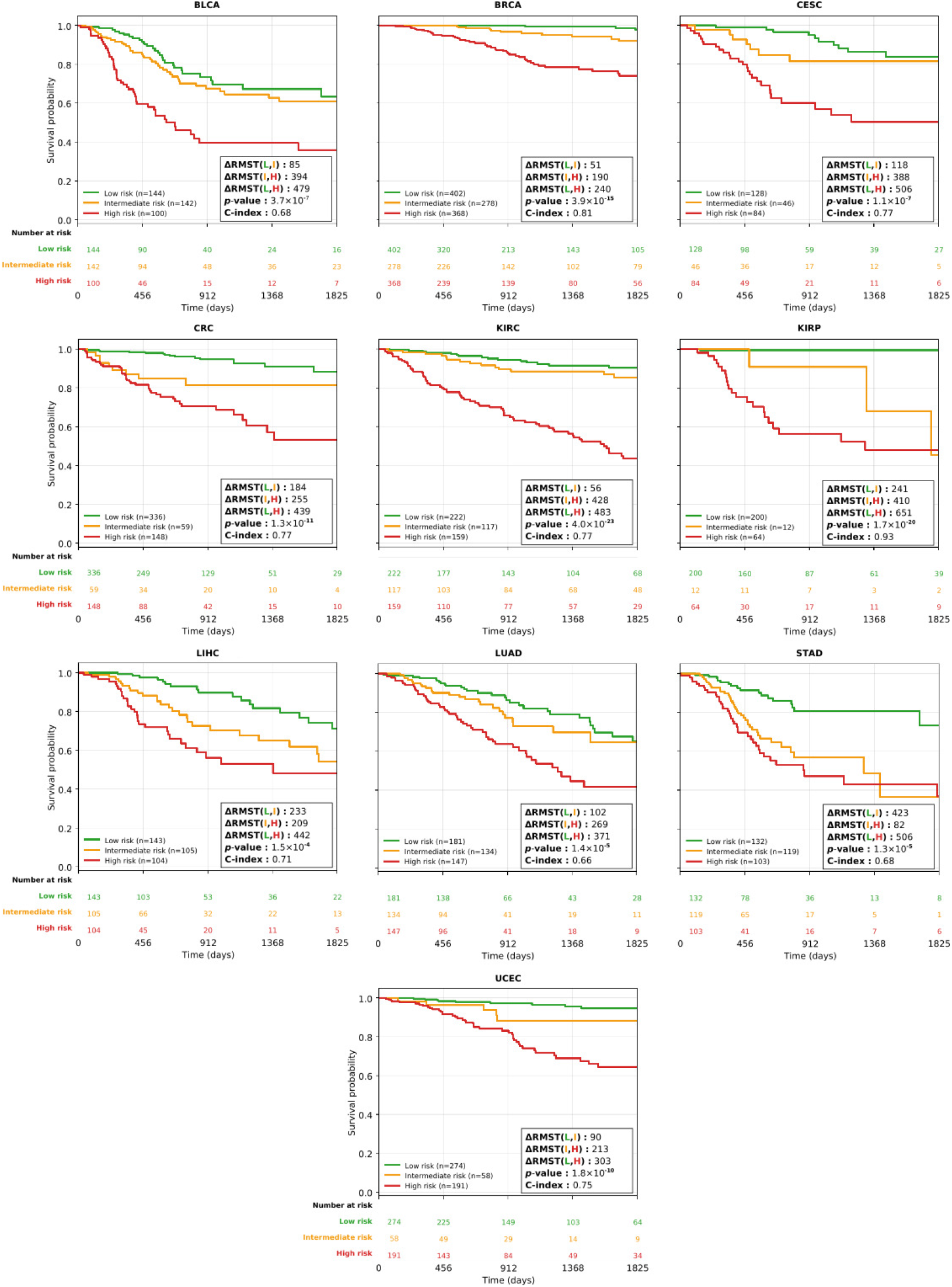
Kaplan-Meier curves of disease-specific survival for the risk groups discovered by GOTSurv for the gene expression cohorts, obtained from the consensus assignment across bootstrap resamples. Each panel reports the difference in restricted mean survival time (ΔRMST) between risk groups, the multivariate log-rank *p*-value, and the concordance index. Group sizes are given in the legend, and the number of patients at risk is shown beneath each panel.

We next compared GOTSurv with established survival models to assess whether directly learnt groups provide stronger stratification than post-hoc discretisation. GOTSurv generally achieved larger adjacent-group RMST differences (Fig. 3A) and stronger log-rank separation (Table 1). For CRC gene expression, it produced the largest RMST difference and a median *p*-value of 0.002, compared with 0.007 for CoxPH and 0.024 for SSVM, while GBM and RSF were non-significant. This pattern was broadly consistent across cohorts; CoxPH was the strongest baseline but less consistently significant. For WSIs, cohorts where GOTSurv was non-significant also showed limited separation across the baselines.

**Table 1.**
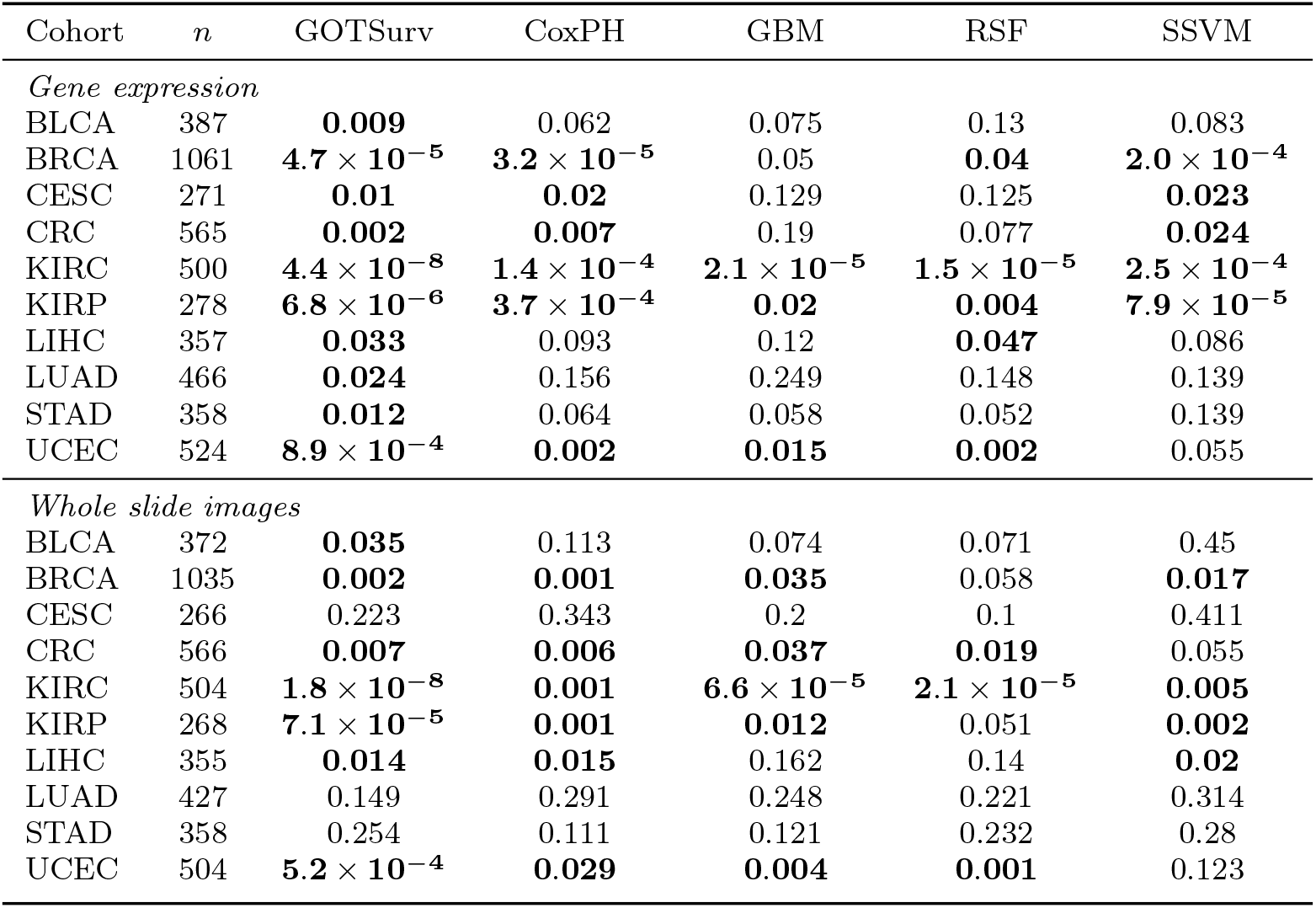
Cohort sizes and median multivariate log-rank *p*-values across 100 bootstrap resamples. Values below 0.05 are shown in bold. For the baseline models, risk groups were obtained by post-hoc tertile discretisation of the predicted risk scores, whereas GOTSurv groups were taken directly from the grouping head.

**Fig. 3.**
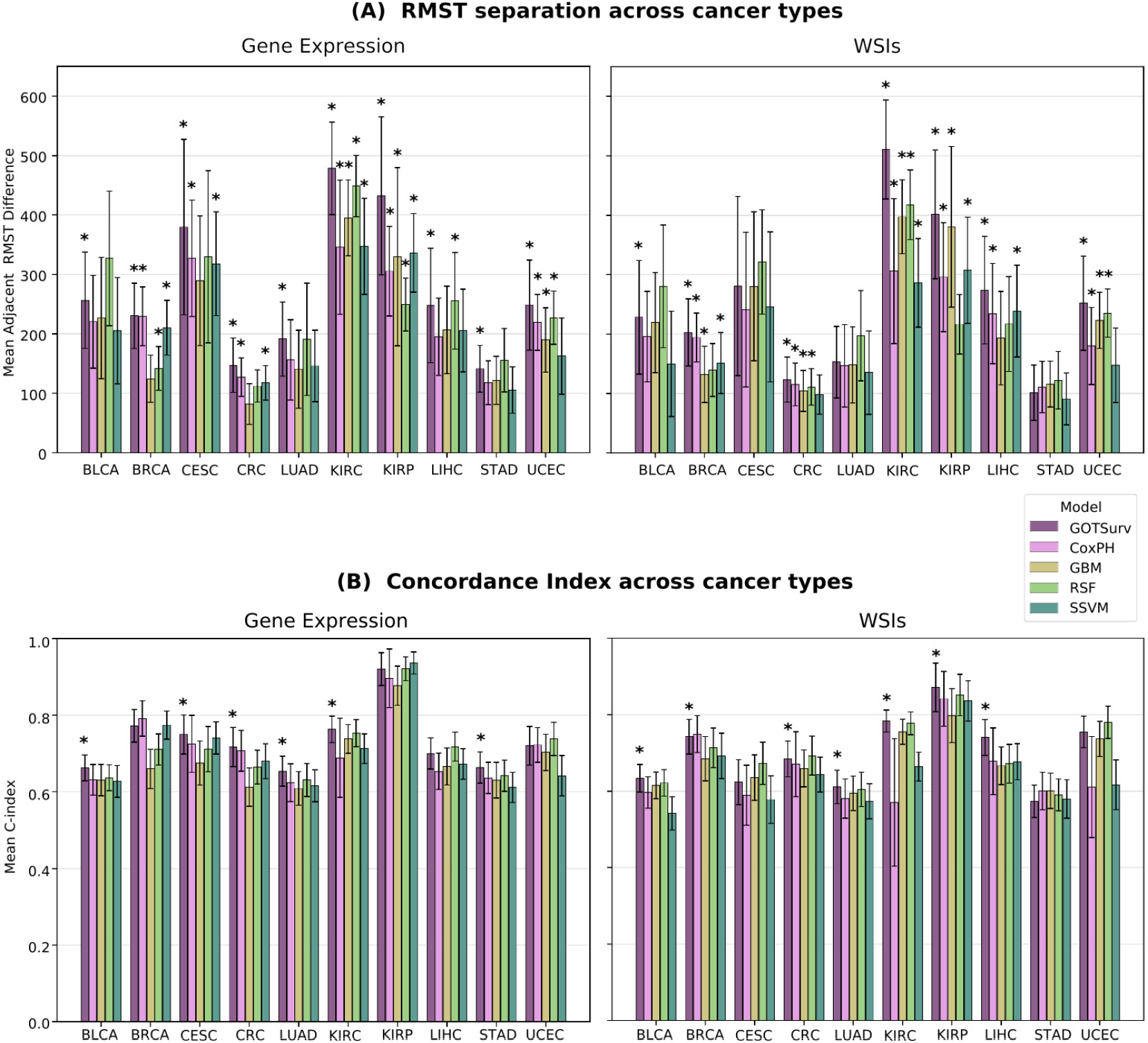
Risk stratification and risk prediction performance of GOTSurv and baseline models across ten cancer types and two modalities. Bars show the mean over 100 bootstrap resamples, with error bars denoting one standard deviation. **(A)** Mean difference in restricted mean survival time (RMST) for gene expression (left) and WSI (right) features, between adjacent risk groups, for the same modalities. For the baseline models, risk groups were obtained by post-hoc tertile discretisation of the predicted risk scores, whereas GOTSurv groups were taken directly from the grouping head. Asterisks in (B) denote statistically significant separation across the three risk groups, defined as a median multivariate log-rank *p <* 0.05 across the 100 bootstrap resamples. **(B)** Mean concordance index (C-index), asterisks denote comparable or better performance for GOTSurv against the best performing baseline.

We next compared C-index performance across models. GOTSurv was generally competitive with the strongest baselines, either achieving the highest mean C-index or showing no significant difference from the best-performing model in many cohorts (Fig. 3B). For example, on CRC WSI data, GOTSurv achieved a mean C-index of 0.685, compared with 0.693 for RSF, with no significant difference across matched bootstrap runs. Thus, the stronger stratification achieved by GOTSurv generally did not come at the expense of survival-ranking performance.

### 3.2 GOTSurv Reveals Patient-Specific Progression Trajectories

Having established that the learnt risk groups represent distinct survival states, we next examine progression at the individual-patient level as population summaries can obscure molecular differences. GOTSurv maps each sample to a patient-specific transport-derived target in the neighbouring higher-risk group, allowing patients within the same group to follow different progression trajectories. Fig. 4 shows selected BRCA patients from each risk group in a Partial Least Squares (PLS) 2D projection of the feature space, each alongside its barycentric target and the corresponding shift in gene-expression profile. Remaining cohort are in Supplementary Fig.S2–S6.

**Fig. 4.**
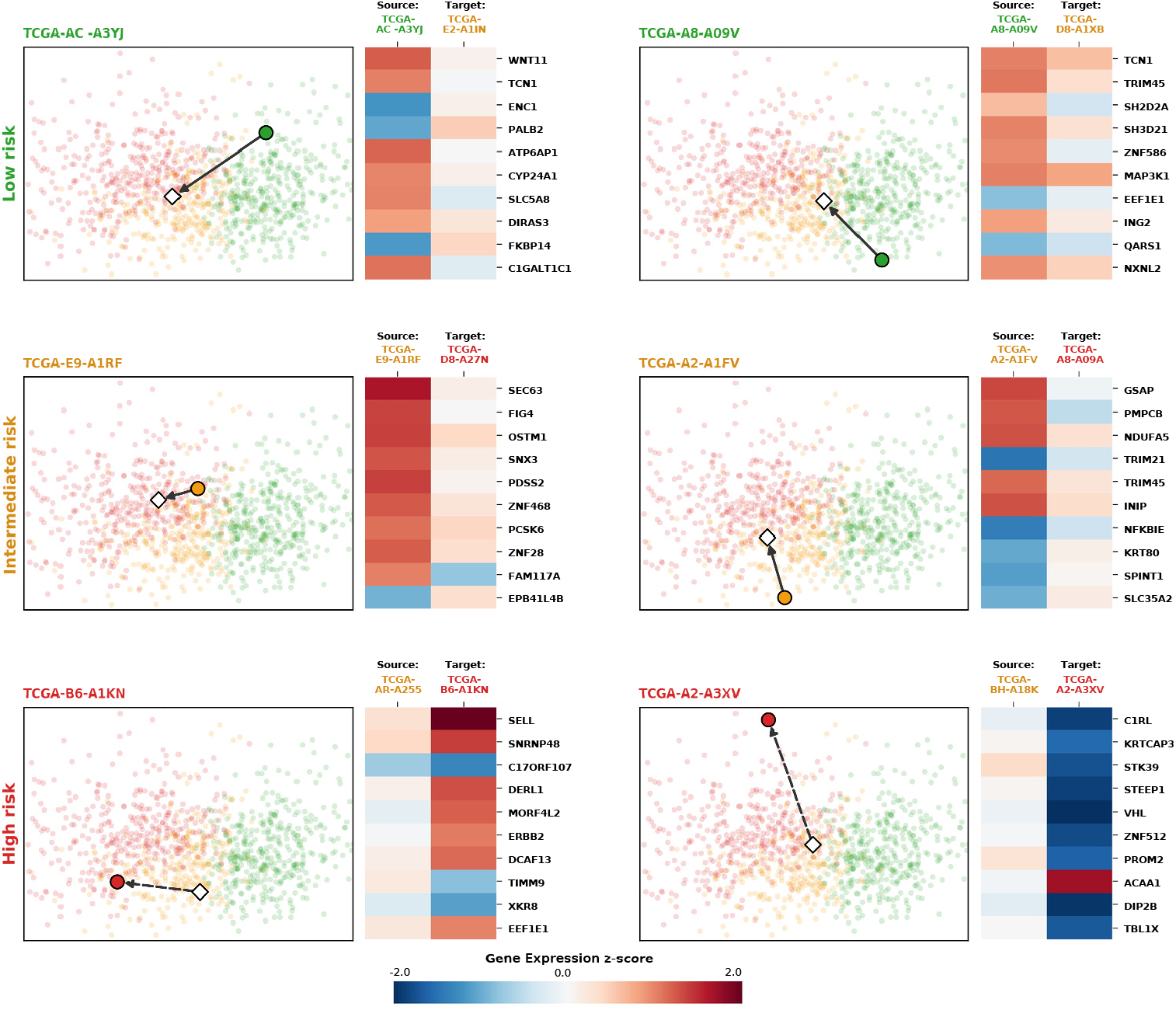
Patient-specific progression profiles in BRCA. Selected patients from each risk group are shown in the PLS feature space together with their transport-derived barycentric target. The heatmaps compare the gene-expression profile of each patient with its corresponding sample in the neighbouring risk group for the features showing the largest changes. For low- and intermediate-risk patients, the inferred target lies in the next higher-risk group, while high-risk patients are compared with their inferred predecessor in the intermediate-risk group.

Patients within the same risk group were mapped to different barycentric targets, showing that similar predicted risk can correspond to distinct inferred progression routes. These targets also differed molecularly from their source patients. For example, the low-risk patient showed higher *TCN1* expression than its inferred intermediate-risk target, consistent with reports linking high *TCN1* to favourable BRCA prognosis and reduced proliferation and invasion [32]. Conversely, the high-risk patient showed lower *KRTCAP3* expression than its inferred intermediate-risk predecessor, consistent with its reported association with unfavourable BRCA prognosis [21]. These patient-specific differences may capture progression features missed by group-level summaries. The same analysis applies to WSIs, where each patient is linked to an inferred slide target (Supplementary Fig. S7–S11).

While the individual profiles capture patient-specific variation, aggregating the progression vectors provides a complementary view of the overall direction of change across the cohort. Fig. 5 projects the learnt whitened latent space into two dimensions using PLS and overlays the progression flow as arrows. The risk gradient is expected because the PLS projection is supervised by predicted risk, whereas the progression vectors are derived independently from the transport couplings. Despite this, the vectors generally point towards higher-risk regions while following locally varying directions, indicating multiple progression routes rather than a single shared trajectory. The overall geometry differs across cancer types, but the transport remains broadly aligned with increasing risk while preserving cohort-specific progression patterns.

**Fig. 5.**
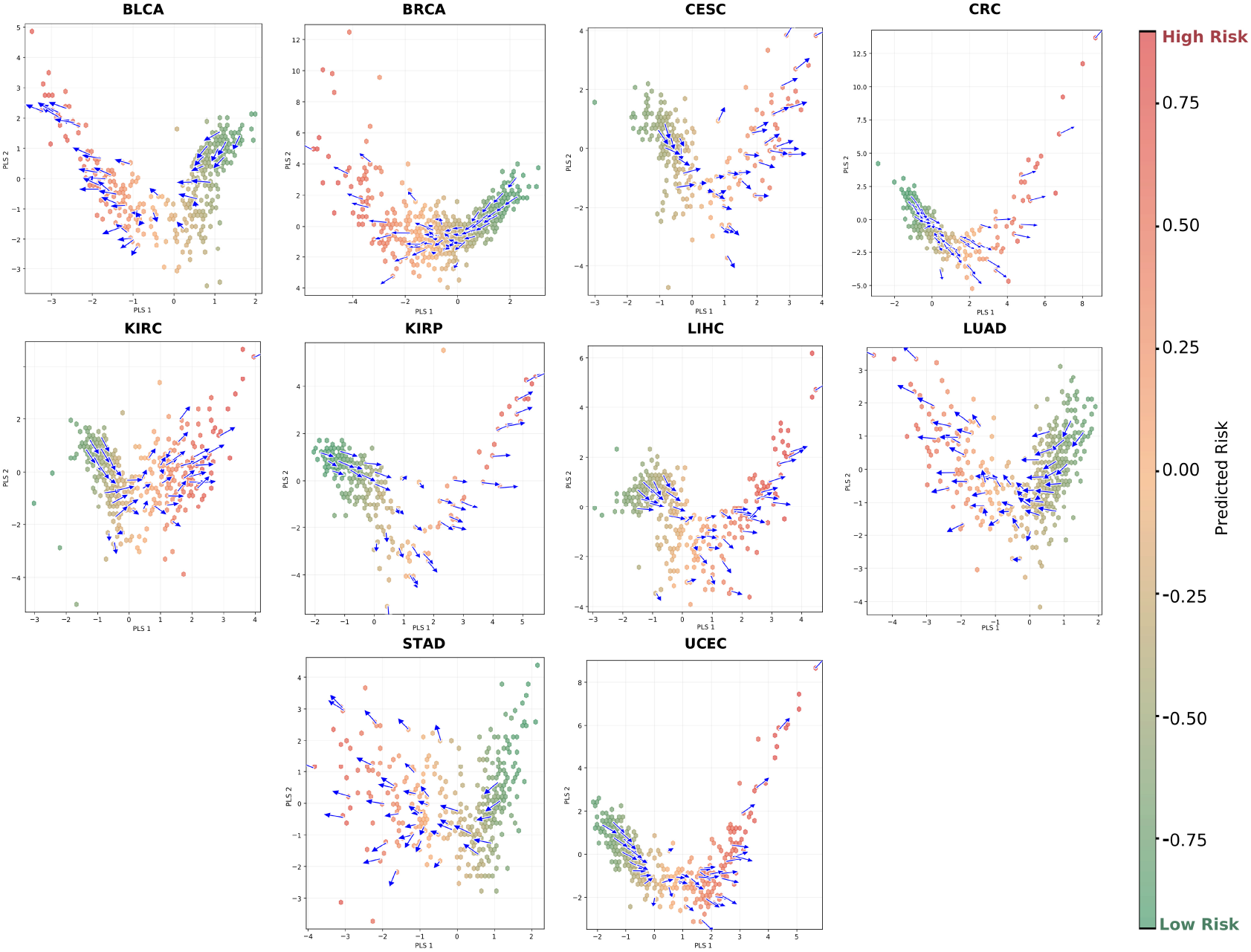
Learnt risk trajectories inferred by GOTSurv for the gene expression cohorts. Samples are projected into two dimensions from the learnt latent space and coloured by predicted risk, using green for low risk, orange for intermediate risk, and red for high risk. Arrows indicate average OT-derived drift vectors from each risk group towards the adjacent higher-risk group.

### 3.3 Risk Groups Capture Distinct Molecular Progression States

To determine whether the learnt risk groups correspond to distinct molecular states, we compared gene expression across the three groups. We identified the most differentially expressed genes using a Kruskal–Wallis test with Benjamini–Hochberg FDR correction, and visualised their expression from low to high risk for BRCA and CRC (Fig. 6), with the remaining cohorts shown in Supplementary Fig. S12.

**Fig. 6.**
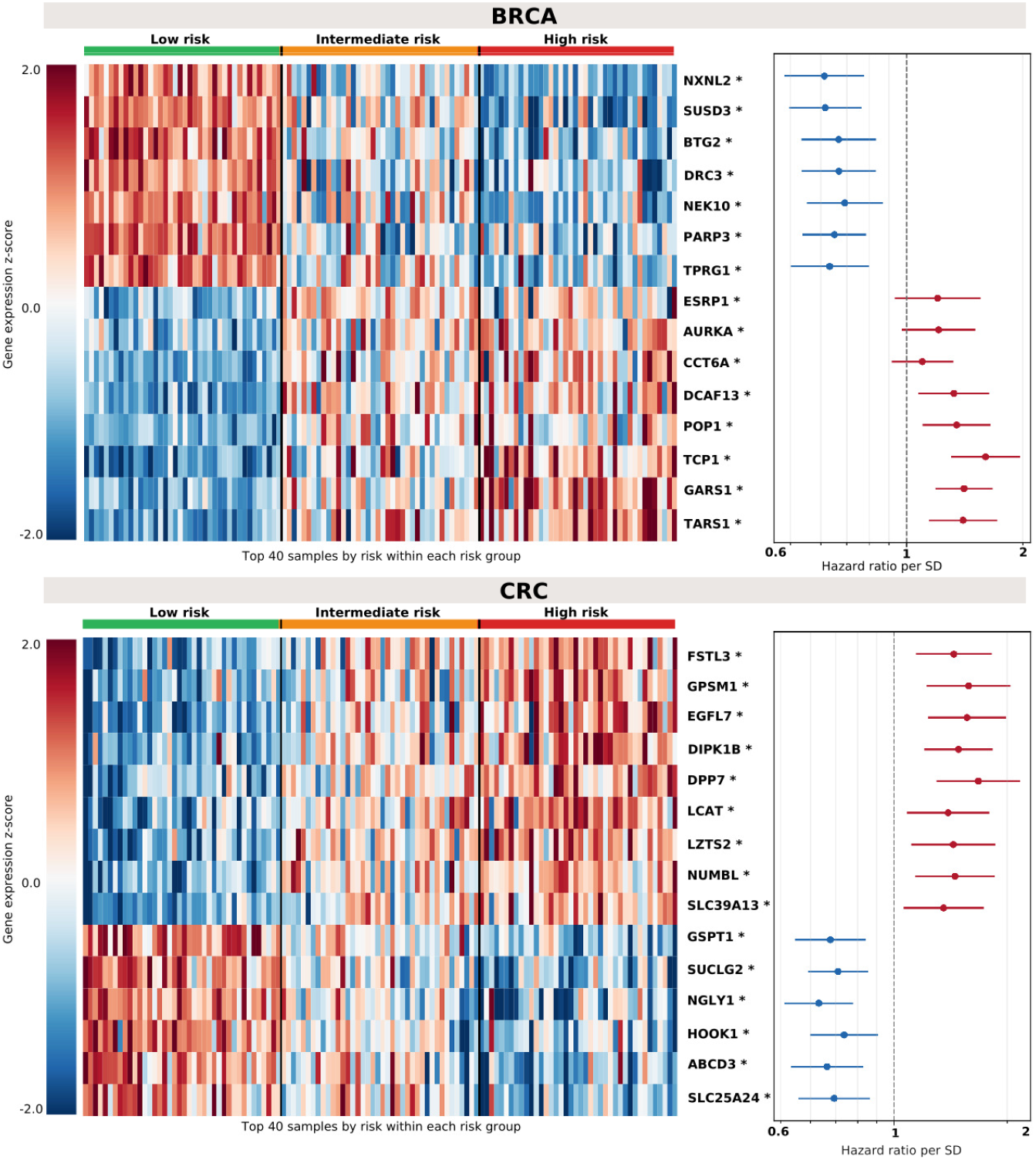
Molecular characterisation of the GOTSurv risk groups for BRCA (top) and CRC (bottom). Heatmaps show the expression (*z*-score) of the genes most differentially expressed across the three risk groups (Kruskal-Wallis, Benjamini-Hochberg FDR; asterisks mark *q <* 0.001), with the 40 highestrisk samples per group ordered from low to high risk. Forest plots show the corresponding univariate Cox hazard ratios per standard deviation increase in gene expression, with 95% confidence intervals.

In both cohorts, the most differentially expressed genes shifted progressively across low-, intermediate-, and high-risk groups rather than distinguishing only the extremes. Some genes decreased with increasing risk, while others showed the opposite pattern. For instance, in BRCA, *NXNL2* decreased towards the high-risk group, consistent with its reported favourable prognostic association [33], while in CRC, *FSTL3* increased with risk and has been shown to promote tumour progression through HIF1 signalling [34].

To test whether these expression differences are prognostically coherent, we related each gene to survival directly through a univariate Cox model. The resulting hazard ratios per standard deviation agree with the heatmap in every case. Genes elevated in the high-risk group carry hazard ratios above one, and genes elevated in the low-risk group carry hazard ratios below one. The groups therefore differ along a molecular axis whose direction matches independent survival associations, indicating that the stratification reflects prognostically meaningful biology rather than an arbitrary partition of expression.

### 3.4 Distinct Progression Patterns Exist Within Risk Groups

Identifying a group of patients within the same risk stratum who follow different biological progression routes can reveal heterogeneity that is not captured by risk stratification alone. To focus specifically on variation beyond predicted risk, we residualised the gene expression progression vectors with respect to the risk score, removing the component explained by risk. Within each risk group, we then clustered the residualised progression vectors into *s* = 2 subgroups using *k*-means. This allows us to identify groups of patients with comparable risk but distinct inferred progression patterns.

For BRCA and CRC, the resulting subgroups were molecularly distinct within each risk level (Fig. 7), with the remaining cohorts shown in Supplementary Fig. S13. Within the intermediate-risk BRCA group, Subgroup 2 showed higher expression of *GZMB, HLA-DOB*, and *SPIB*, whereas Subgroup 1 showed an opposing pattern. The elevated expression of *GZMB, HLA-DOB*, and *SPIB* is consistent with a more immune-active phenotype, characterised by stronger cytotoxic lymphocyte activity and B-cell-related antigen presentation [35, 36]. In CRC, the low-risk subgroups were similarly distinct, with Subgroup 1 showing higher expression of genes including *OMA1* and *BCL10*, which have been linked to hypoxia-driven metabolic reprogramming and NF-*κ*B signalling [37, 38].

**Fig. 7.**
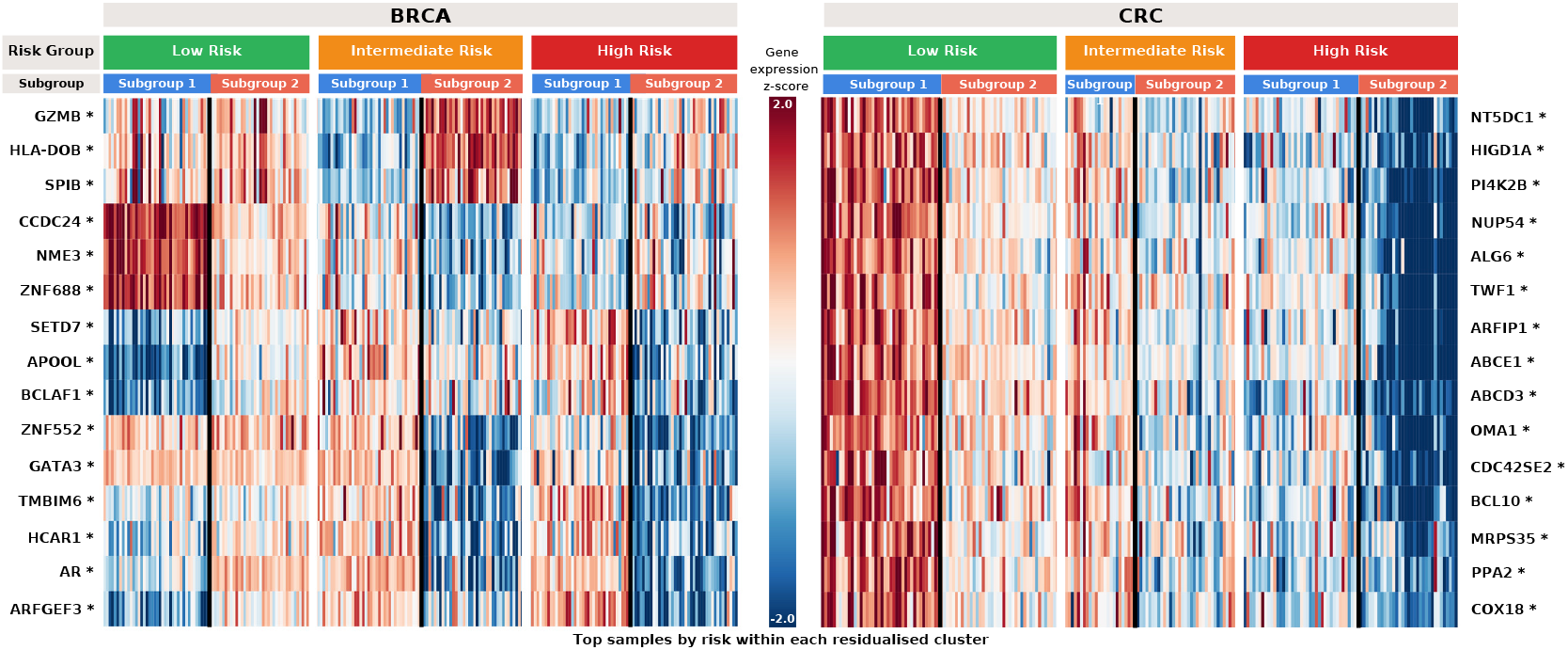
Within-group heterogeneity of the GOTSurv risk groups for BRCA (left) and CRC (right). Within each risk group, samples are split into two subgroups obtained from risk-residualised progression vectors, and heatmaps show the expression (*z*-score) of the genes best separating them (Mann-Whitney, Benjamini-Hochberg FDR; asterisks mark *q <* 0.05). Subgroup blocks are ordered by expression for display only.

We next asked whether the molecular differences between progression subgroups reflected broader biological programmes. Genes were ranked by the magnitude and direction of their transport-derived changes and analysed using preranked GSEA with Hallmark, Reactome, and literature-derived gene sets. A positive Normalised Enrichment Scores (NES) means a gene set concentrated among the genes that increase (the top of the ranking), indicating a gene-set up-regulated along progression. A negative NES identifies a set concentrated among the genes that decrease (the bottom of the ranking), indicating a gene-set down-regulated along progression. The resulting enrichment profiles for CRC are shown in Fig. 8, with remaining cohorts in Supplementary Fig.S14.

**Fig. 8.**
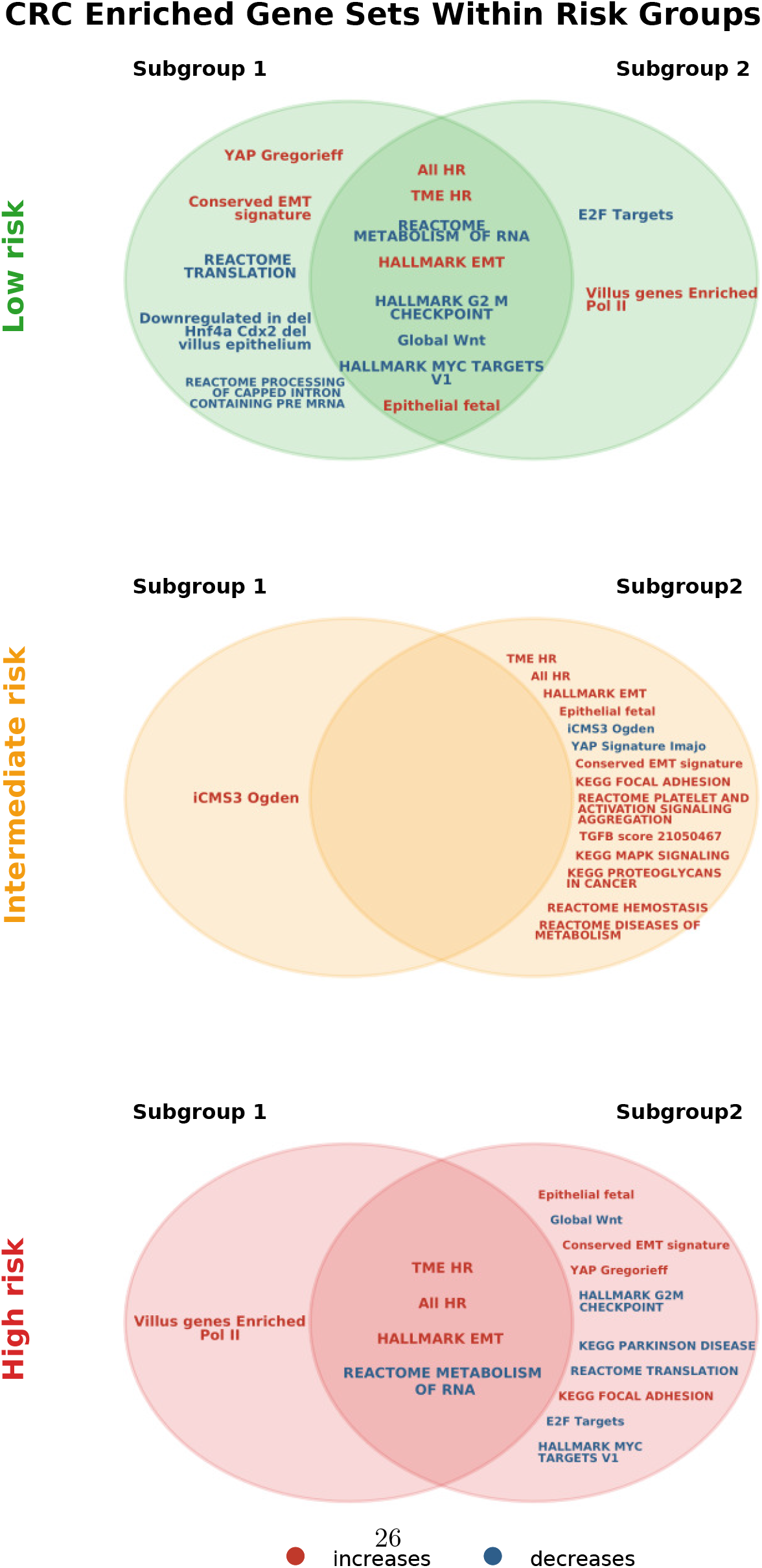
Venn diagram showing shared and subgroup-specific progression-associated gene sets within the CRC risk groups. Shared regions indicate gene sets enriched in both progression subgroups, while non-overlapping regions indicate subgroup-specific biological programmes.

The low- and high-risk subgroups showed substantial overlap in their progression-associated biological programmes, suggesting that patients at these risk levels may share a common underlying route of progression despite subgroup-specific differences. In contrast, the intermediate-risk subgroups were markedly more divergent. In particular, one subgroup showed increasing EMT- and tumour-microenvironment-associated programmes together with a decrease in iCMS3, whereas the other showed the opposite iCMS3 direction. Since iCMS3 represents an intrinsic epithelial state in CRC, while EMT and stromal programmes reflect a mesenchymal acquisition and distinct microevironmental axis [39, 40], this pattern is consistent with alternative progression routes at intermediate risk, including a shift away from an iCMS3-like epithelial programme towards a more mesenchymal and microenvironment-associated state [39–41].

Together, these results show that patients within the same risk group can follow distinct inferred progression patterns, characterised by differences at both the gene and broader molecular-programme levels.

### 3.5 Risk Ranking and Stratification Generalises to External Cohorts

To assess whether GOTSurv transfers beyond the cohorts it was developed on, we evaluated it on two independent datasets, ABCTB for WSIs and the CPTAC colorectal cohort for gene expression. Performance results are shown in Table 2.

**Table 2.**
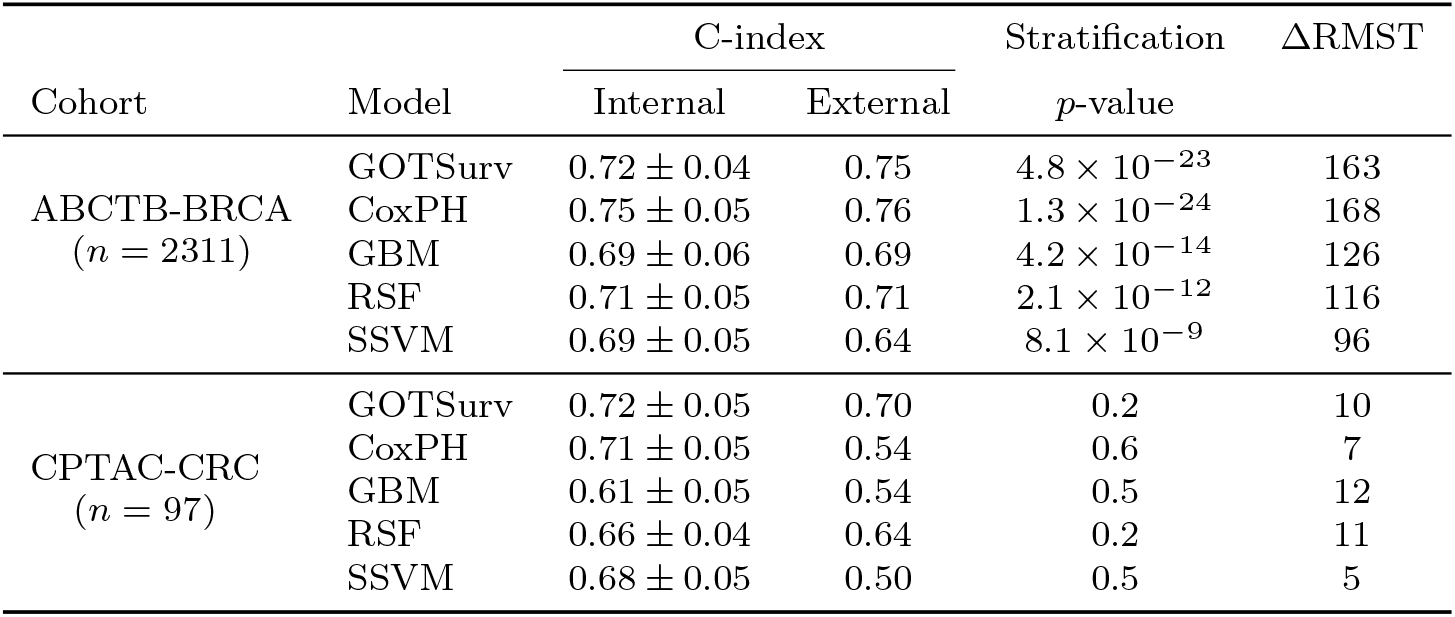
External validation of risk ranking and stratification. Models trained on TCGA are applied to independent cohorts without refitting. Internal C-index values are reported as mean*±*SD across bootstrap resamples, while external metrics are evaluated on the independent cohort. RMST denotes the mean adjacent-group restricted mean survival time difference.

| Cohort | Model | C-index |  | Stratification | ΔRMST |
|---|---|---|---|---|---|
|  |  | Internal | External | *p*-value |  |
| ABCTB-BRCA<br>(*n* = 2311) | GOTSurv | 0.72 ± 0.04 | 0.75 | 4.8 × 10<sup>−23</sup> | 163 |
|  | CoxPH | 0.75 ± 0.05 | 0.76 | 1.3 × 10<sup>−24</sup> | 168 |
|  | GBM | 0.69 ± 0.06 | 0.69 | 4.2 × 10<sup>−14</sup> | 126 |
|  | RSF | 0.71 ± 0.05 | 0.71 | 2.1 × 10<sup>−12</sup> | 116 |
|  | SSVM | 0.69 ± 0.05 | 0.64 | 8.1 × 10<sup>−9</sup> | 96 |
| CPTAC-CRC<br>(*n* = 97) | GOTSurv | 0.72 ± 0.05 | 0.70 | 0.2 | 10 |
|  | CoxPH | 0.71 ± 0.05 | 0.54 | 0.6 | 7 |
|  | GBM | 0.61 ± 0.05 | 0.54 | 0.5 | 12 |
|  | RSF | 0.66 ± 0.04 | 0.64 | 0.2 | 11 |
|  | SSVM | 0.68 ± 0.05 | 0.50 | 0.5 | 5 |

In ABCTB, ranking performance transferred well across models. GOTSurv achieved an external C-index of 0.75, comparable to CoxPH at 0.76, and produced strong risk-group separation (*p* = 4.8 × 10^−23^) with a RMST difference of 163 days.

CoxPH showed similar stratification in this cohort, with *p* = 1.3 × 10^−24^ and a RMST difference of 168 days, while the remaining baselines showed smaller risk separation. Transfer to CPTAC was more challenging. GOTSurv largely retained its internal ranking performance and achieved the highest external C-index, whereas some baselines showed larger declines, such as CoxPH from 0.71 to 0.54. However, no model produced significant risk-group separation in CPTAC, indicating that external validation in this cohort primarily supports transfer of ranking performance rather than stratification.

Overall, these results show that GOTSurv generalises to independent cohorts, retaining both ranking performance and automatic stratification in the WSI cohort, and preserving ranking accuracy in the gene-expression cohort where several baselines showed substantial declines.

### 3.6 Abstention Analysis

To assess whether uncertain assignments affect performance, we performed confidence-based abstention across 100 bootstrap resamples, ranking patients by the assignment confidence *c*_*i*_ and progressively excluding the lowest-confidence samples. At each abstention rate, RMST, multivariate log-rank *p*-value, and C-index were recomputed and compared with random abstention. Differences from the full-cohort baseline were assessed using the Wilcoxon signed-rank test across matched resamples. BRCA and CRC results for the gene expression cohorts are shown in Fig. 9 and Table 3, with remaining cohorts in Supplementary Fig. S15.

**Table 3.**
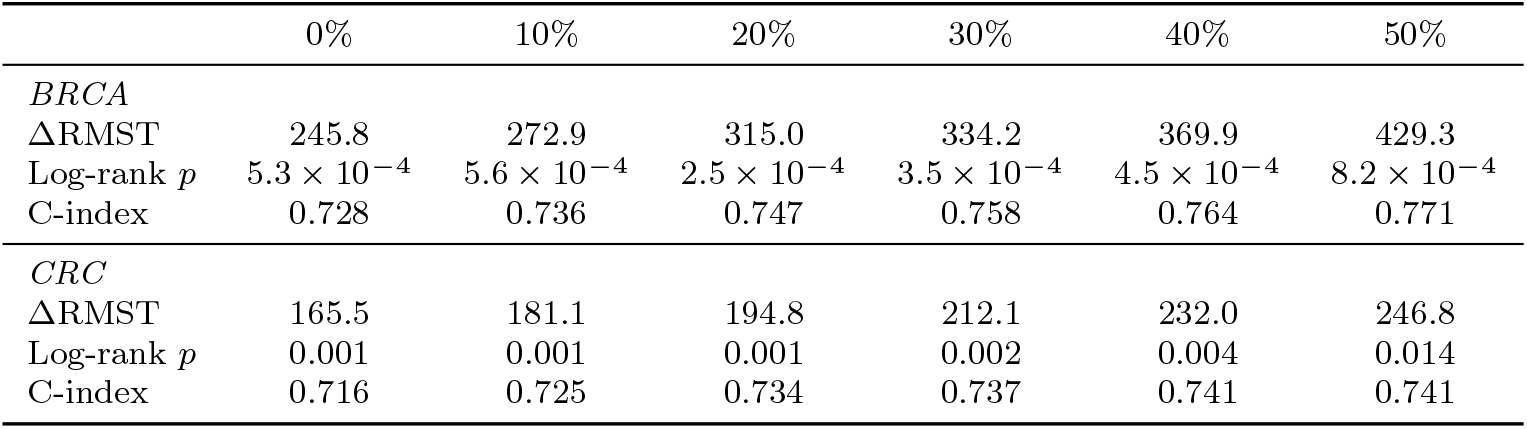
Risk stratification metrics under confidence-based abstention. Values are aggregated across bootstrap resamples, with C-index and adjacent ΔRMST reported as means and log-rank *p* as the median.

|  | 0% | 10% | 20% | 30% | 40% | 50% |
|---|---|---|---|---|---|---|
| *BRCA* |  |  |  |  |  |  |
| ΔRMST | 245.8 | 272.9 | 315.0 | 334.2 | 369.9 | 429.3 |
| Log-rank *p* | 5.3 × 10<sup>−4</sup> | 5.6 × 10<sup>−4</sup> | 2.5 × 10<sup>−4</sup> | 3.5 × 10<sup>−4</sup> | 4.5 × 10<sup>−4</sup> | 8.2 × 10<sup>−4</sup> |
| C-index | 0.728 | 0.736 | 0.747 | 0.758 | 0.764 | 0.771 |
| *CRC* |  |  |  |  |  |  |
| ΔRMST | 165.5 | 181.1 | 194.8 | 212.1 | 232.0 | 246.8 |
| Log-rank *p* | 0.001 | 0.001 | 0.001 | 0.002 | 0.004 | 0.014 |
| C-index | 0.716 | 0.725 | 0.734 | 0.737 | 0.741 | 0.741 |

**Fig. 9.**
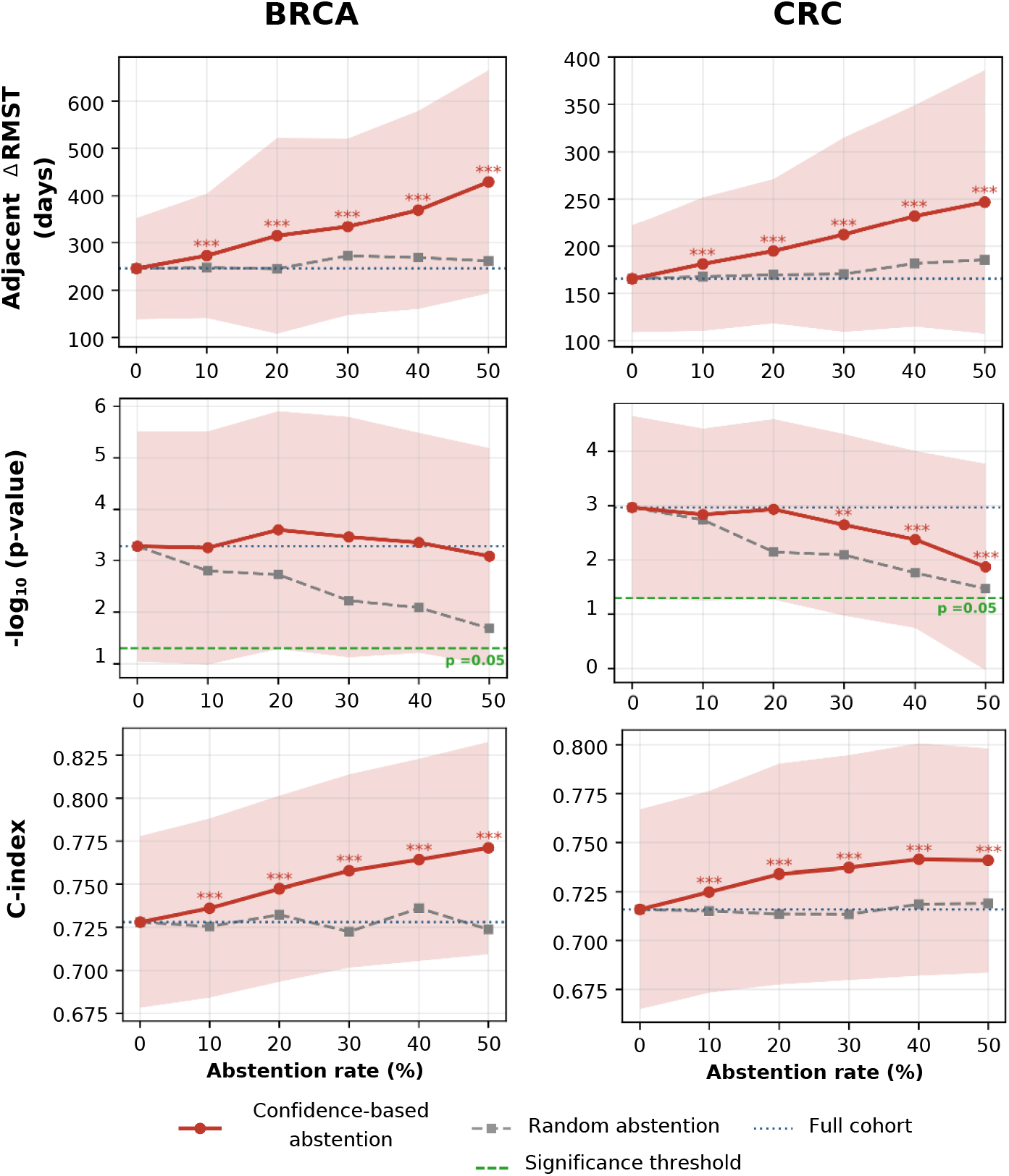
Confidence-based abstention analysis for BRCA and CRC. Changes in adjacent-group RMST separation, multivariate log-rank significance, and C-index as the abstention rate increases. Confidence-based abstention is shown against random abstention and full-cohort performance. Shaded regions show variability across resamples, and asterisks indicate significant differences from random abstention. Asterisks indicate a significant difference from the full-cohort (0% abstention) baseline using the Wilcoxon signed-rank test across matched bootstrap resamples.

As lower-confidence patients were progressively excluded, survival separation improved in both cohorts. In BRCA, the mean adjacent-group RMST difference increased from 246 days at baseline to 429 days at 50% abstention, while in CRC it increased from 165 to 247 days. Despite retaining only the more confidently assigned patients, the risk groups remained significantly separated, with log-rank *p*-values below 0.05 at 50% abstention in both BRCA (*p* = 8.2 × 10^−4^) and CRC (*p* = 0.014). Ranking performance also improved, with the C-index increasing from 0.72 up to 0.77 in BRCA and from 0.72 up to 0.74 in CRC. At higher abstention rates (≥ 30%), however, performance variability increased as fewer patients were retained, highlighting the need to balance the benefits of excluding low-confidence assignments against the loss of sample size.

We further assessed confidence-based abstention on the external validation cohorts (Table 4). In ABCTB, abstention improved both stratification and ranking, with adjacent RMST increasing from 163 to 424 days and C-index from 0.749 to 0.807 before a slight decline at the highest abstention rate. In CPTAC, finer abstention increments increased RMST from 10 to 17 days while C-index remained stable, although risk-group separation remained non-significant throughout.

**Table 4.** Confidence-based abstention on the independent external validation cohorts. GOTSurv was trained on the corresponding TCGA cohort and applied to the external cohort without refitting. Values are reported on the retained external samples at each abstention rate.

| *ABCTB-BRCA* |  |  |  |  |  |  |
| Abstention rate | 0% | 10% | 20% | 30% | 40% | 50% |
|---|---|---|---|---|---|---|
| ΔRMST | 163 | 175 | 188 | 207 | 265 | 424 |
| Log-rank *p* | 4.8 × 10<sup>−23</sup> | 6.7 × 10<sup>−26</sup> | 2.0 × 10<sup>−26</sup> | 8.4 × 10<sup>−26</sup> | 2.5 × 10<sup>−23</sup> | 6.5 × 10<sup>−20</sup> |
| C-index | 0.749 | 0.764 | 0.781 | 0.793 | 0.807 | 0.795 |
| *CPTAC-CRC* |  |  |  |  |  |  |
| Abstention rate | 0% | 2% | 4% | 6% | 8% | 10% |
| ΔRMST | 10 | 12 | 13 | 14 | 15 | 17 |
| Log-rank *p* | 0.236 | 0.232 | 0.240 | 0.230 | 0.218 | 0.204 |
| C-index | 0.695 | 0.696 | 0.693 | 0.690 | 0.691 | 0.695 |

Overall, *c*_*i*_ provided a useful measure of assignment confidence, with abstention generally preserving or improving performance among retained patients.

## 4 Discussion

We introduced GOTSurv to infer patient-specific progression trajectories towards higher survival risk from cross-sectional data. Unlike population-level analyses, GOT-Surv resolves progression at the level of individual patients, revealing feature changes that would otherwise be averaged across groups. The learnt risk groups represented distinct survival and molecular states, providing a meaningful structure over which progression could be modelled. Within these risk levels, GOTSurv further revealed subgroups of patients following different progression patterns, reflected in distinct molecular profiles and biological programmes. Across ten cancer cohorts and two modalities, GOTSurv produced stronger risk stratification while maintaining competitive survival-ranking performance, with these findings also supported by two independent external cohorts.

These results highlight the main contribution of GOTSurv in resolving progression at the level of individual patients rather than assigning a single pattern of change to an entire risk group. Each sample is linked to its own set of inferred targets in the neighbouring group, producing a patient-specific progression profile. This differs from standard differential expression analysis, which summarises changes at the group level. The results showed that this distinction was important, as patients within the same risk group were often linked to different inferred targets and followed different inferred routes towards higher risk. A single group-level summary therefore captures only part of the progression structure and can obscure substantial variation between individuals. Future work could explore learning the number of risk groups adaptively from the data rather than specifying *K* in advance. Additional experiments on larger external cohorts and further data modalities, as well as multimodal integration, could further assess and extend the applicability of GOTSurv.

## Supplementary information

The supplementary material contains the ablation study and its results table, together with the figures for the remaining cancer types and modalities referenced in the text. A separate excel file is included for the full comparison results.

## Acknowledgements

EA is sponsored by the Saudi Cultural Bureau in London, UK. The authors acknowledge support from the Tissue Image Analysis Centre at the University of Warwick. We thank the Australian Breast Cancer Tissue Bank (ABCTB) for providing H&E images, and especially appreciate the assistance of Mythily Sachchithananthan and Judith Heads at the ABCTB. Tissues and samples were received from the Australian Breast Cancer Tissue Bank which is generously supported by the National Health and Medical Research Council of Australia, The Cancer Institute NSW and the National Breast Cancer Foundation. We are grateful to Asa Ben-Hur (Colorado State University) for helpful input during the early conceptualisation of the grouping loss. We also would like to thank Aimée Seltan (KU Leuven) for expert input on the CRC gene-signature findings.

## Declarations

### 4.2 Author Contributions

E.A. and F.M. contributed to the conceptualisation of the study. E.A. performed the formal analysis and visualization, and led the development of methodology in collaboration with F.M. and G.W. Supervision was provided by F.M. E.A. drafted the original manuscript, and all authors contributed to the review and editing of the final version.

### 4.3 Data availability

The TCGA data analysed in this study are publicly available through the Genomic Data Commons (https://portal.gdc.cancer.gov). Clinical outcome annotations were taken from the TCGA Pan-Cancer Clinical Data Resource [20]. The slide-level embeddings were derived from the TCGA-UT-8K collection released with TITAN [22]. The CPTAC colorectal cancer data are publicly available through cBioPortal (https://www.cbioportal.org) and are described in [23]. Data and material from the Australian Breast Cancer Tissue Bank (ABCTB) [24] are available to approved researchers on application through the ABCTB (https://abctb.org.au); they were used here under an approved project and cannot be redistributed by the authors.

### 4.4 Code availability

The implementation of GOTSurv, including training scripts, evaluation pipelines, and instructions for reproducing the results in this study, is available at: https://github.com/EtharZaid/GOTSurv.

