## Supplementary File for "Discovering Prognostic Risk Strata and Heterogeneous Progression Routes from Cross-Sectional Survival Data"

### 1 Derivation of loss gradients

For completeness, we derive the two complementary gradients of the ordering objective. Let

$$A = \sum_{(u,v)\in\mathcal{P}} a_{uv}, \qquad H = \sum_{(u,v)\in\mathcal{P}} a_{uv}h_{uv}, \qquad \mathcal{L}_{\text{ord}} = \frac{H}{A}.$$

Assuming $A > 0$ and holding the ranking penalties fixed, the quotient rule gives

$$\begin{aligned}
\frac{\partial \mathcal{L}_{\text{ord}}}{\partial a_{ij}} &= \frac{\partial}{\partial a_{ij}}\left(\frac{H}{A}\right) \\
&= \frac{A\frac{\partial H}{\partial a_{ij}} - H\frac{\partial A}{\partial a_{ij}}}{A^2} \\
&= \frac{Ah_{ij} - H}{A^2} \\
&= \frac{h_{ij} - H/A}{A} \\
&= \frac{h_{ij} - \mathbb{E}_{\pi}[h]}{A}.
\end{aligned}$$

Similarly, holding the agreement probabilities fixed,

$$\begin{aligned}
\frac{\partial \mathcal{L}_{\text{ord}}}{\partial h_{ij}} &= \frac{\partial}{\partial h_{ij}}\left(\frac{H}{A}\right) \\
&= \frac{A\frac{\partial H}{\partial h_{ij}} - H\frac{\partial A}{\partial h_{ij}}}{A^2} \\
&= \frac{Aa_{ij} - H(0)}{A^2} \\
&= \frac{a_{ij}}{A} \\
&= \pi_{ij}.
\end{aligned}$$

These derivations treat $a_{ij}$ and $h_{ij}$ as independent intermediate variables. During training, both are determined by the jointly optimised neural-network parameters, so their actual updates reflect contributions from multiple comparable pairs.

### 2 Sensitivity to Optimal Transport Parameters

We assessed the sensitivity of GOTSurv to the two OT parameters controlling entropic smoothing and mass preservation on the BRCA gene expression cohort. We varied blur $\in \{0.1, 0.3, 0.5, 0.7, 1.0\}$ and reach $\in \{0.001, 0.01, 0.1, 1.0\}$, re-running the progression-modelling stage for each combination while keeping the learnt risk-stratification model fixed. Each setting was compared with the parameters used throughout the main experiments (blur $= 0.5$, reach $= 0.01$).

The inferred patient-specific progression directions were highly robust across the parameter grid, with a mean cosine similarity of at least 0.96 relative to the reference setting for every combination. In other words, for a given patient, the inferred progression vector continued to point towards essentially the same region of feature space despite changes in the transport parameters.

### 3 Ablation Study

To assess the contribution of the main components of GOTSurv, we performed a systematic ablation study. The full model was used as the reference setting, with risk groups learnt directly by the model and used to define the progression graph. We compared this with three ablations: replacing the learnt groups with post-hoc quantile-based groups, removing the ranking loss, and removing the proportion penalty. We also report the collapse rate, defined as the percentage of bootstrap runs in which fewer than the specified $K$ risk groups were recovered. Results are shown in Table 1.

The learnt risk groups provided clearer survival stratification than post-hoc quantile grouping. Because both settings use the same continuous risk predictions, their C-index was unchanged. However, the learnt groups achieved stronger survival separation ($p = 6 \times 10^{-4}$ versus $3 \times 10^{-3}$) and a larger RMST gap (234 versus 175 days).

Removing the ranking loss had the largest effect. The C-index decreased from 0.72 to 0.56, risk separation was no longer significant ($p = 0.50$), and the collapse rate increased to 48.8%. This indicates that the ranking objective is important for both prognostic ordering and stable risk-group formation. Removing the proportion penalty had little effect on ranking performance, with the C-index remaining close to the full model ($0.71 \pm 0.05$). However, stratification weakened ($p = 0.04$) and the collapse rate increased to 21.0%. Although the RMST gap increased to 338 days, this occurred

Table 1: Ablation and sensitivity analysis for BRCA. C-index is reported as mean ± standard deviation across bootstrap runs. RMST is the restricted mean survival time gap between adjacent risk groups. Collapse rate denotes the percentage of bootstrap runs in which the model produced fewer than the specified $K$ risk groups.

| **Setting** | **C-index ± SD** | **$p$-value** | **RMST** | **Collapse rate** |
|---|---|---|---|---|
| *Component ablations* | | | | |
| Full model ($K = 3$) | $0.72 \pm 0.04$ | $6 \times 10^{-4}$ | 234 | 0.0% |
| Quantiles | $0.72 \pm 0.04$ | $3 \times 10^{-3}$ | 175 | 0.0% |
| No ranking loss | $0.56 \pm 0.09$ | 0.50 | 185 | 48.8% |
| No proportion loss | $0.71 \pm 0.05$ | 0.04 | 338 | 21.0% |
| *Sensitivity to number of risk groups* | | | | |
| $K = 2$ | $0.72 \pm 0.05$ | $1.2 \times 10^{-3}$ | 334 | 0.0% |
| $K = 4$ | $0.73 \pm 0.05$ | $3.2 \times 10^{-4}$ | 216 | 13.0% |
| $K = 5$ | $0.72 \pm 0.05$ | $4.3 \times 10^{-4}$ | 238 | 34.0% |

alongside less stable group formation and should not be interpreted as improved stratification. The proportion penalty therefore contributes mainly to maintaining stable and well-represented risk groups.

We also assessed sensitivity to the number of risk groups by varying $K$ from 2 to 5. Ranking remained stable across all settings, with C-index values of approximately 0.72-0.73, and all configurations retained significant survival stratification. Stability decreased as $K$ increased, with collapse rates of 13% for $K = 4$ and 34% for $K = 5$. $K = 2$ produced the largest RMST gap but provided only a coarse two-group stratification. $K = 3$ offered a suitable balance between discrimination, survival separation, and stable group formation, supporting its use as the default setting. However, the appropriate $K$ is likely to depend on the cohort, since not all datasets support the same degree of prognostic stratification.

Overall, the ablation study shows that the ranking and proportion terms play complementary roles in predictive performance, stratification, and group stability, while the choice of $K$ controls the level of risk-group resolution supported by the data.

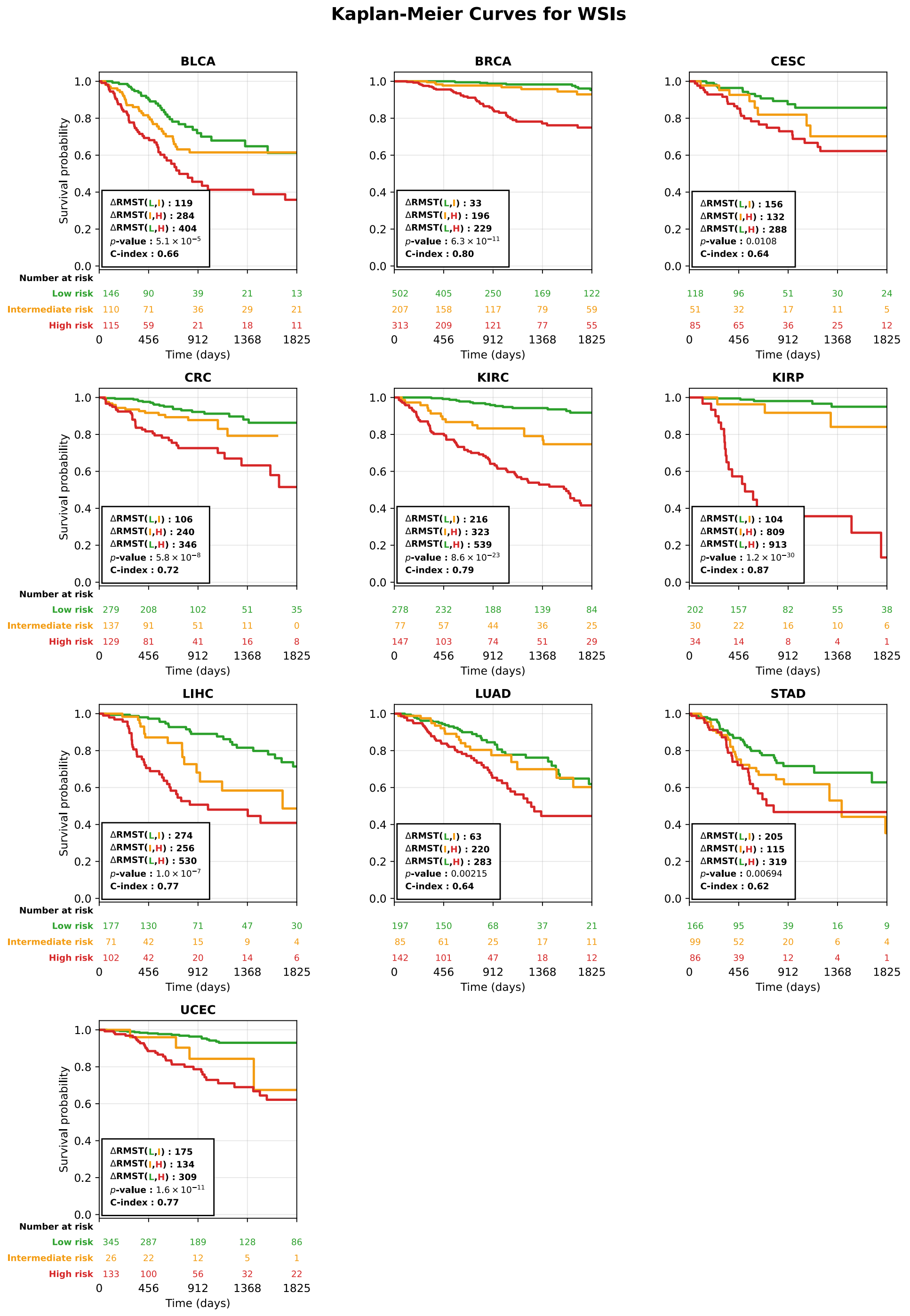


**Fig. S1.** Kaplan–Meier curves of disease-specific survival for the risk groups discovered by GOTSurv for the gene expression cohorts, obtained from the consensus assignment across bootstrap resamples. Each panel reports the concordance index of the consensus risk score, the multivariate log-rank p- value, and the difference in restricted mean survival time (ΔRMST) between adjacent and between extreme risk groups, given as (adjacent, extreme) in days at $\tau = 5$ years. Group sizes are given in the legend, and the number of patients at risk is shown beneath each panel.

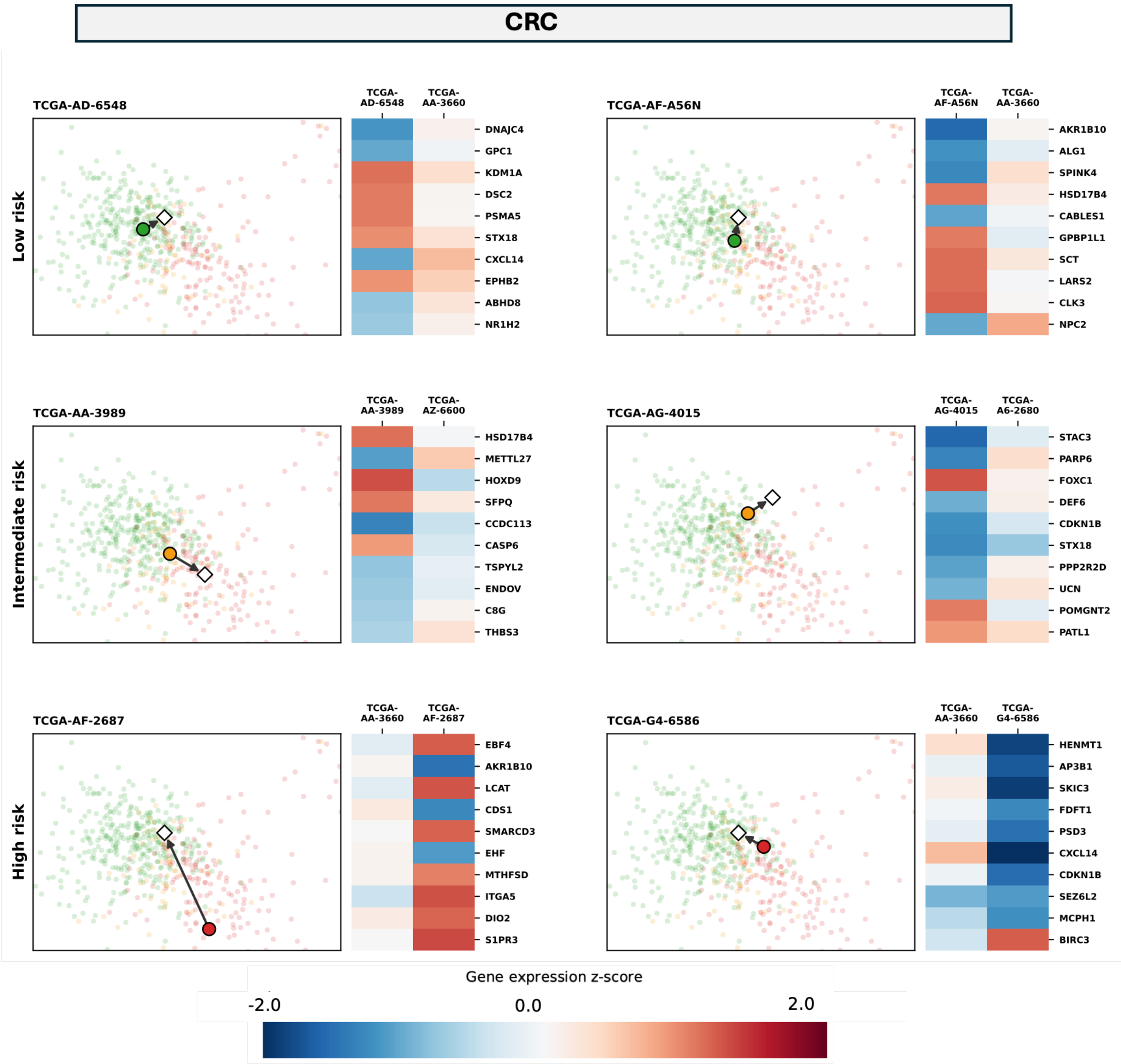


**Fig. S2.** Patient-specific progression profiles CRC. Each panel shows the features changing most for one patient, given as the feature-space progression vector Δx towards its transport-derived target. Red indicates an increase while blue indicates a decrease, and grey bars give the group-level mean shift for the same features. Rows are risk groups, and within each row the three patients are selected at the 10th, 50th and 90th percentiles of cosine similarity to the group-level shift.

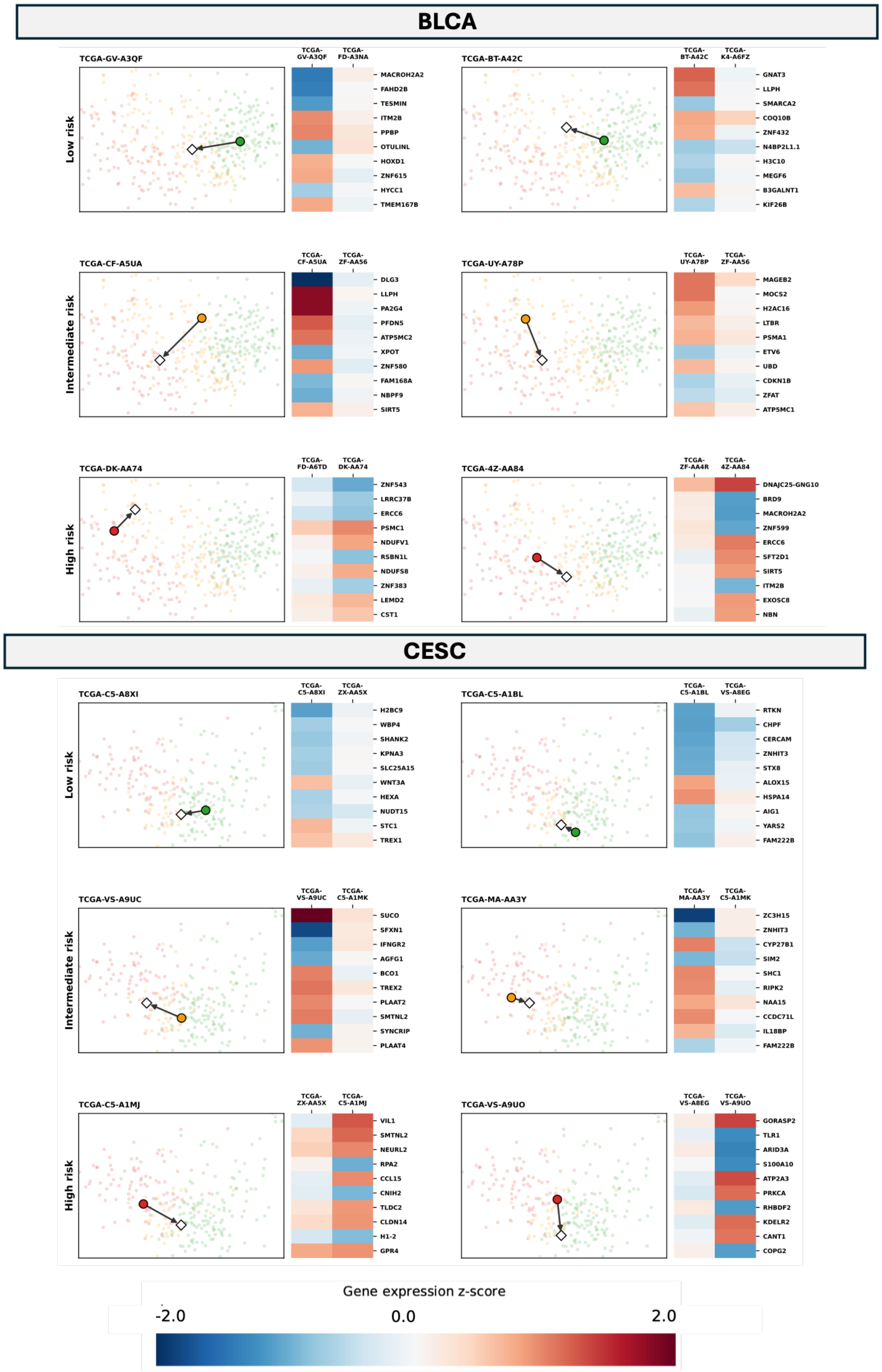


**Fig. S3.** Patient-specific progression profiles in BLCA and CESC. Each panel shows the features changing most for one patient, given as the feature-space progression vector Δx towards its transport-derived target. Red indicates an increase while blue indicates a decrease, and grey bars give the group-level mean shift for the same features. Rows are risk groups, and within each row the three patients are selected at the 10th, 50th, and 90th percentiles of cosine similarity to the group-level shift.

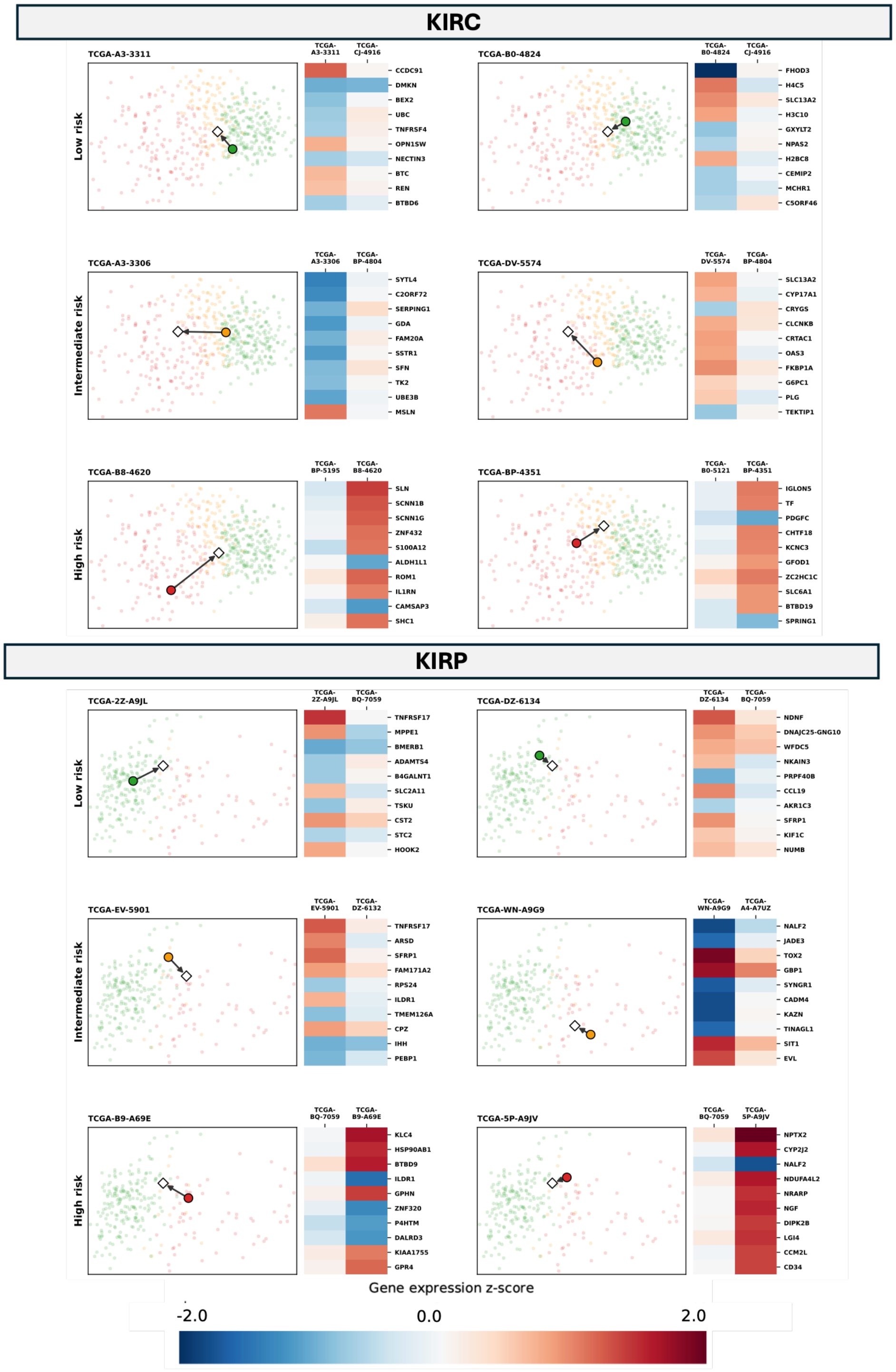


**Fig. S4.** Patient-specific progression profiles in KIRC and KIRP. Each panel shows the features changing most for one patient, given as the feature-space progression vector Δx towards its transport-derived target. Red indicates an increase while blue indicates a decrease, and grey bars give the group-level mean shift for the same features. Rows are risk groups, and within each row the three patients are selected at the 10th, 50th and 90th percentiles of cosine similarity to the group-level shift.

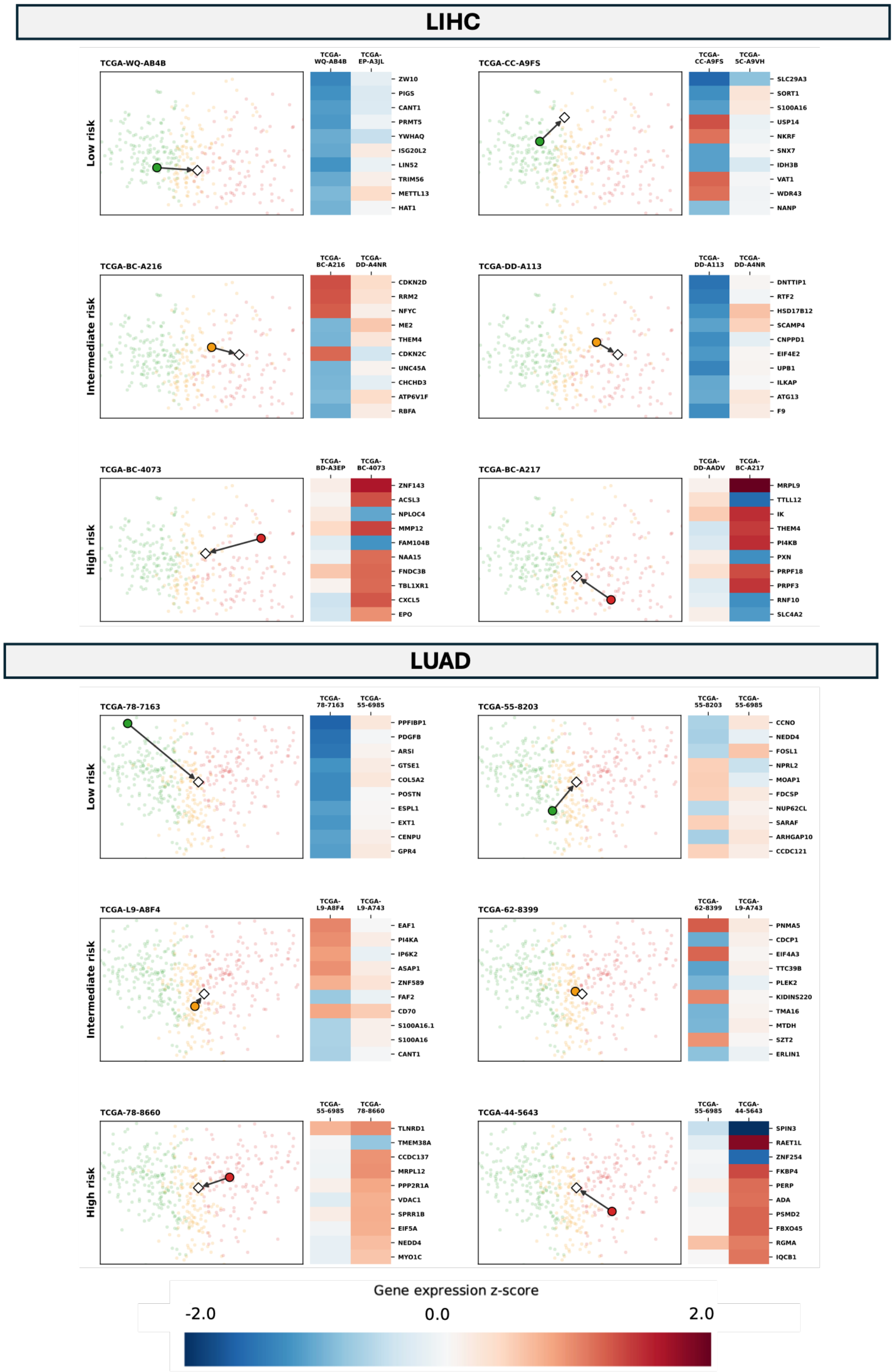


**Fig. S5.** Patient-specific progression profiles in LIHC and LUAD. Each panel shows the features changing most for one patient, given as the feature-space progression vector Δx towards its transport-derived target. Red indicates an increase while blue indicates a decrease, and grey bars give the group-level mean shift for the same features. Rows are risk groups, and within each row the three patients are selected at the 10th, 50th and 90th percentiles of cosine similarity to the group-level shift.

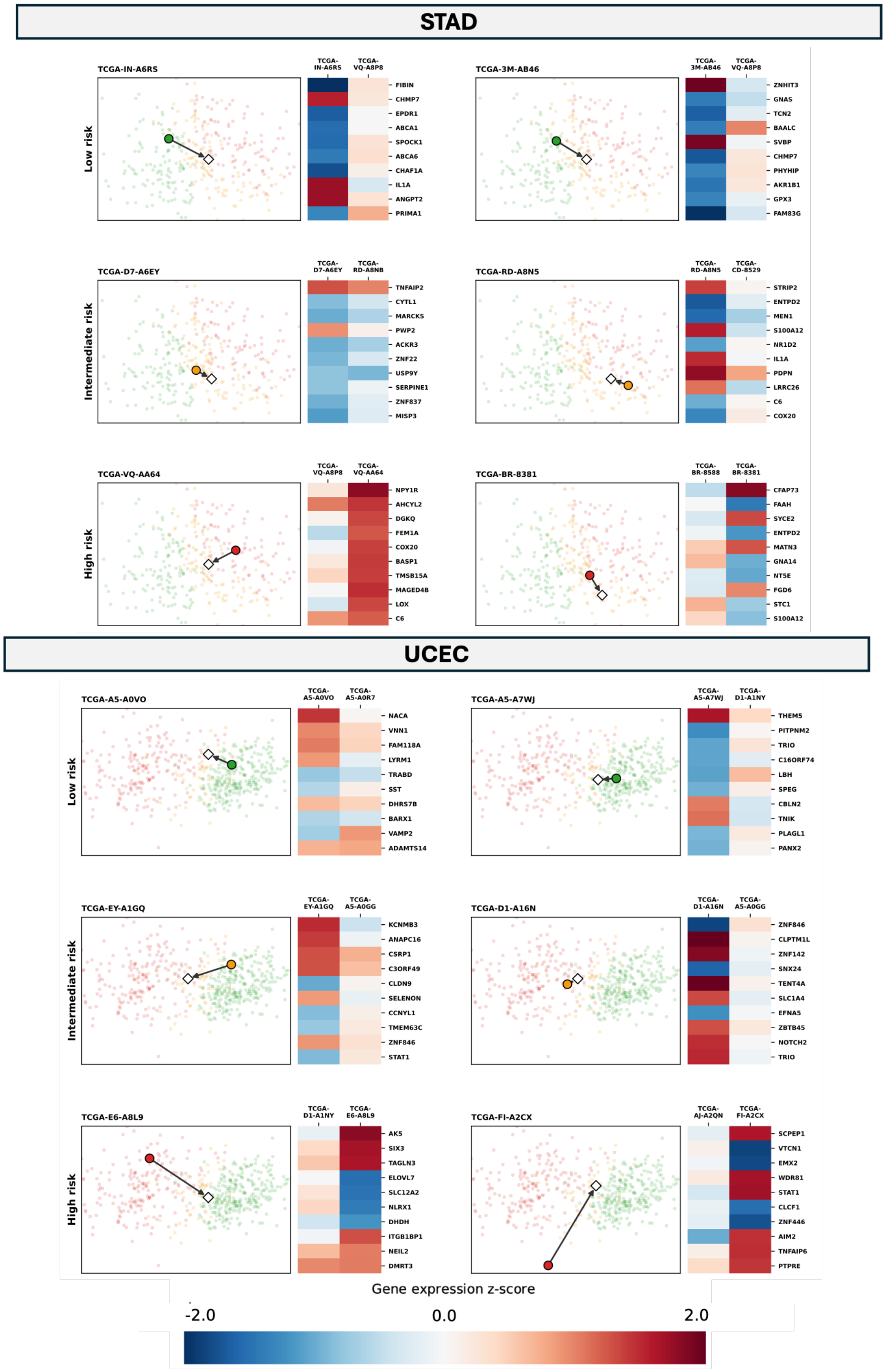


**Fig. S6.** Patient-specific progression profiles in STAD and UCEC. Each panel shows the features changing most for one patient, given as the feature-space progression vector Δx towards its transport-derived target. Red indicates an increase while blue indicates a decrease, and grey bars give the group-level mean shift for the same features. Rows are risk groups, and within each row the three patients are selected at the 10th, 50th and 90th percentiles of cosine similarity to the group-level shift.

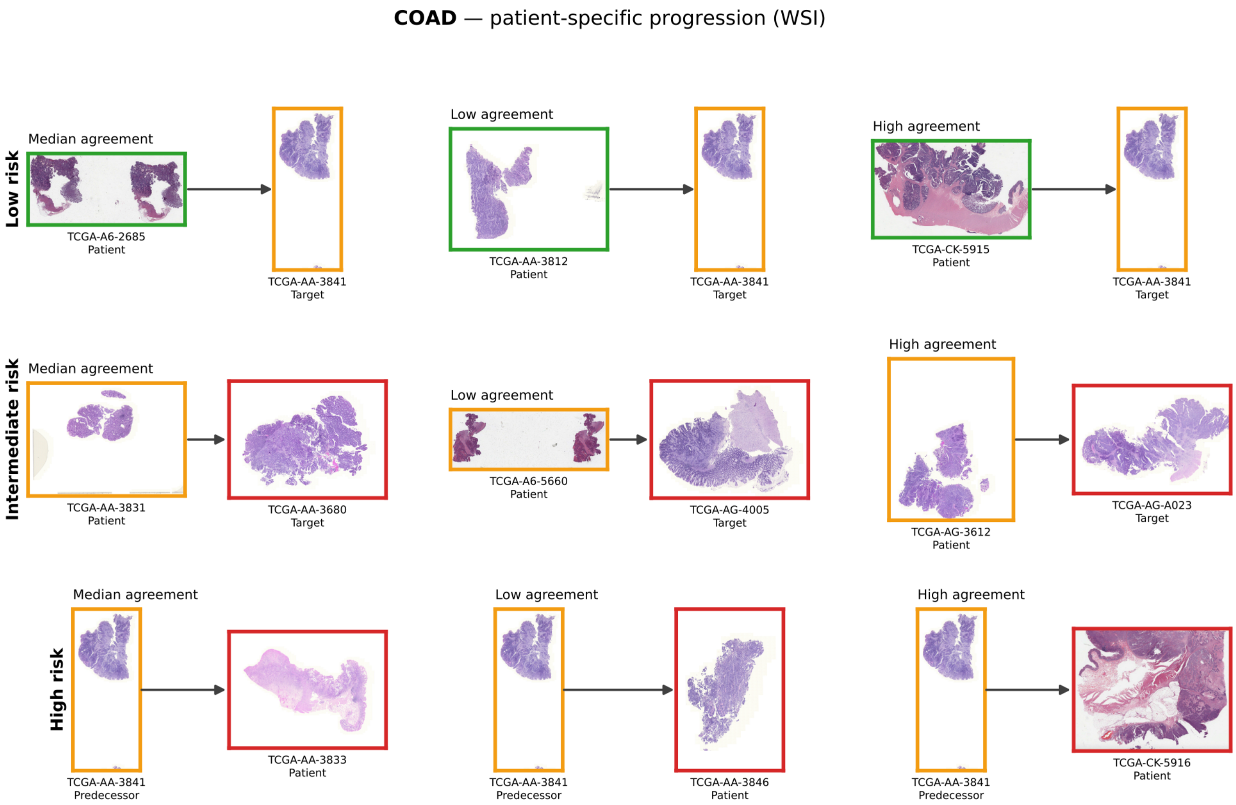


**Fig.S7.** Patient-specific progression in CRC from whole slide images. Selected patients from each risk group are paired with their transport-derived counterpart in the neighbouring risk group. Arrows point towards higher risk, so for high-risk patients the counterpart is the inferred predecessor. Border colours denote risk group.

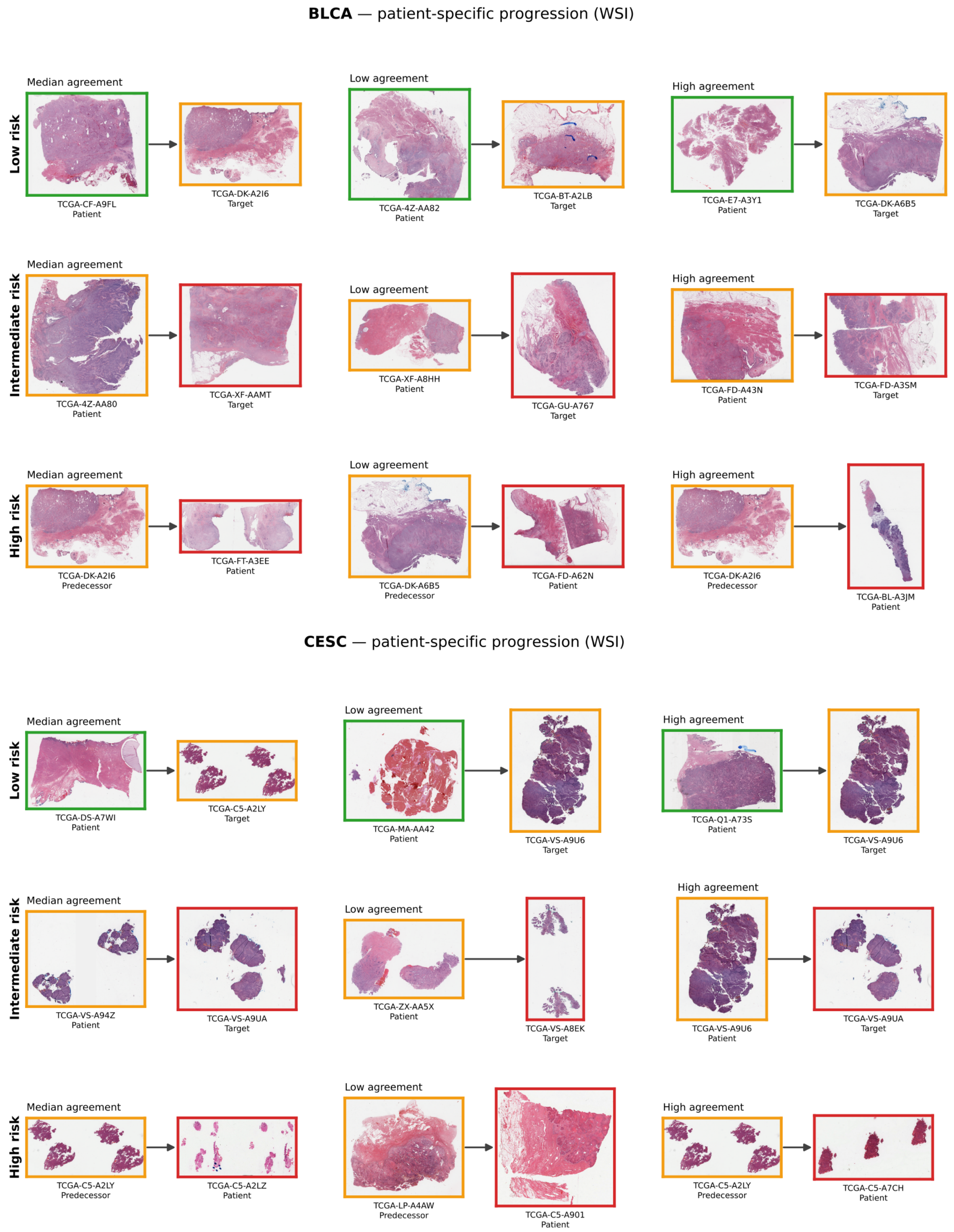


**Fig.S8.** Patient-specific progression in BLCA and CESC from whole slide images. Selected patients from each risk group are paired with their transport-derived counterpart in the neighbouring risk group. Arrows point towards higher risk, so for high-risk patients the counterpart is the inferred predecessor. Border qqlours denote risk group.

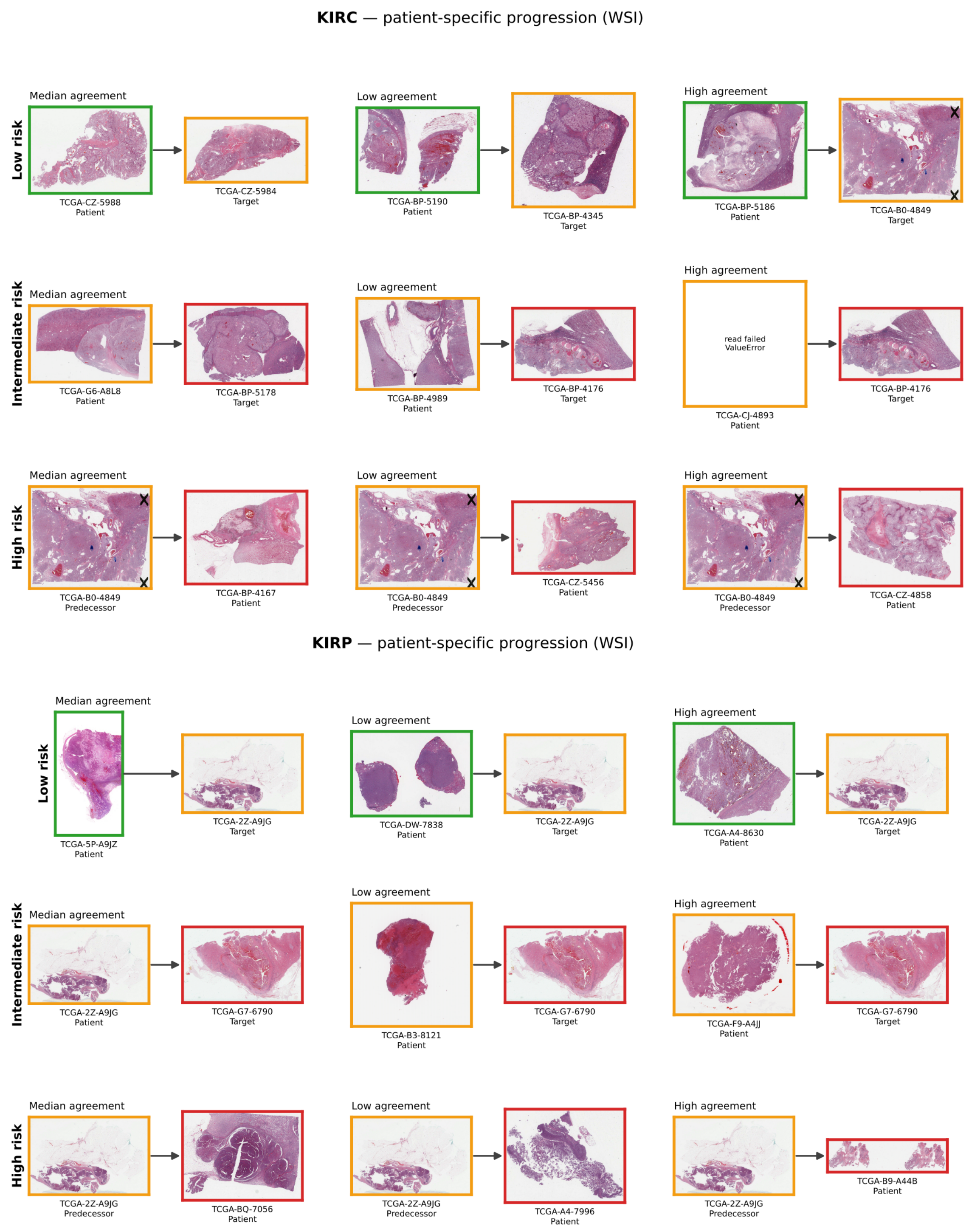


**Fig.S9.** Patient-specific progression in KIRP and KIRC from whole slide images. Selected patients from each risk group are paired with their transport-derived counterpart in the neighbouring risk group. Arrows point towards higher risk, so for high-risk patients the counterpart is the inferred predecessor. Border colours denote risk group.

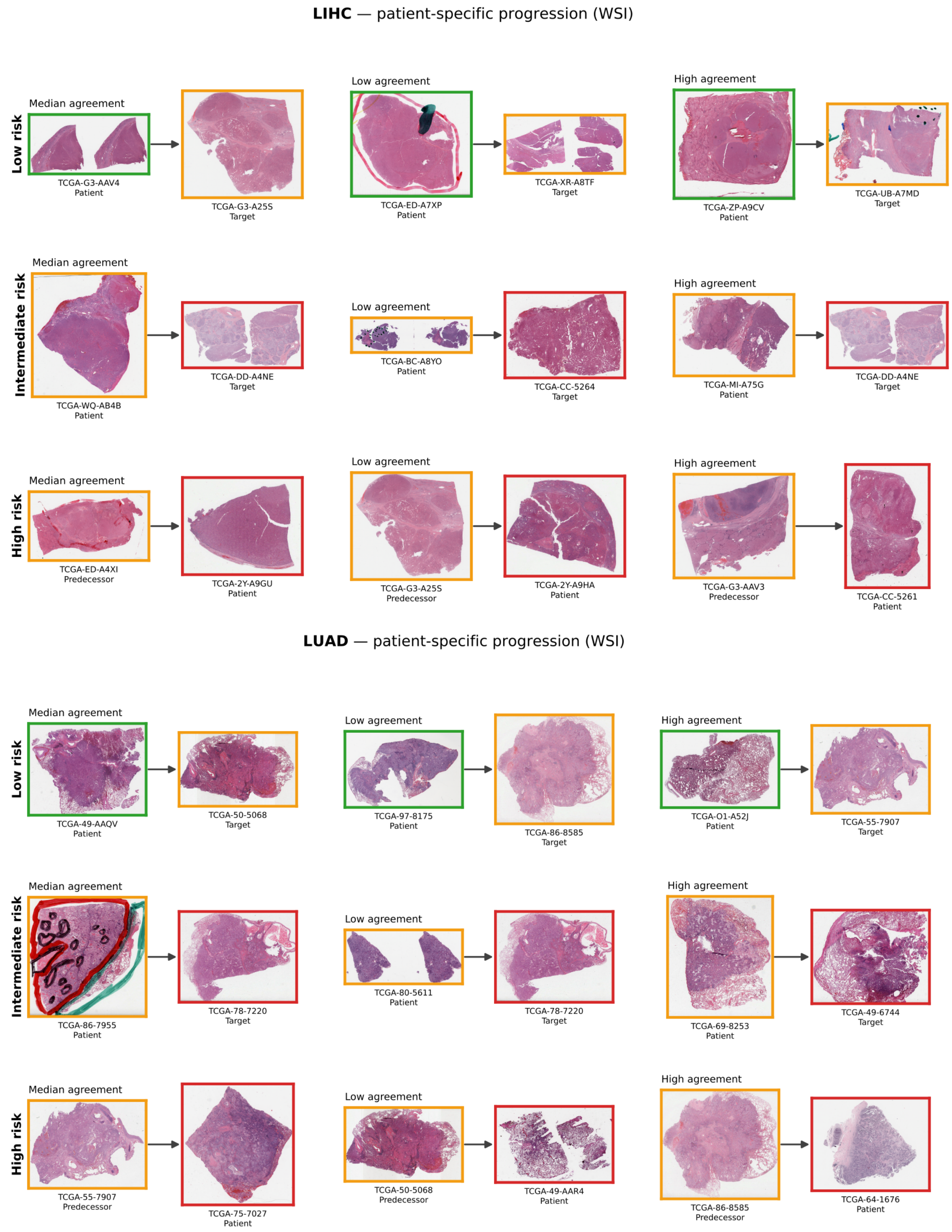


**Fig.S10.** Patient-specific progression in LIHC and LUAD from whole slide images. Selected patients from each risk group are paired with their transport-derived counterpart in the neighbouring risk group. Arrows point towards higher risk, so for high-risk patients the counterpart is the inferred predecessor. Border colours denote risk group.

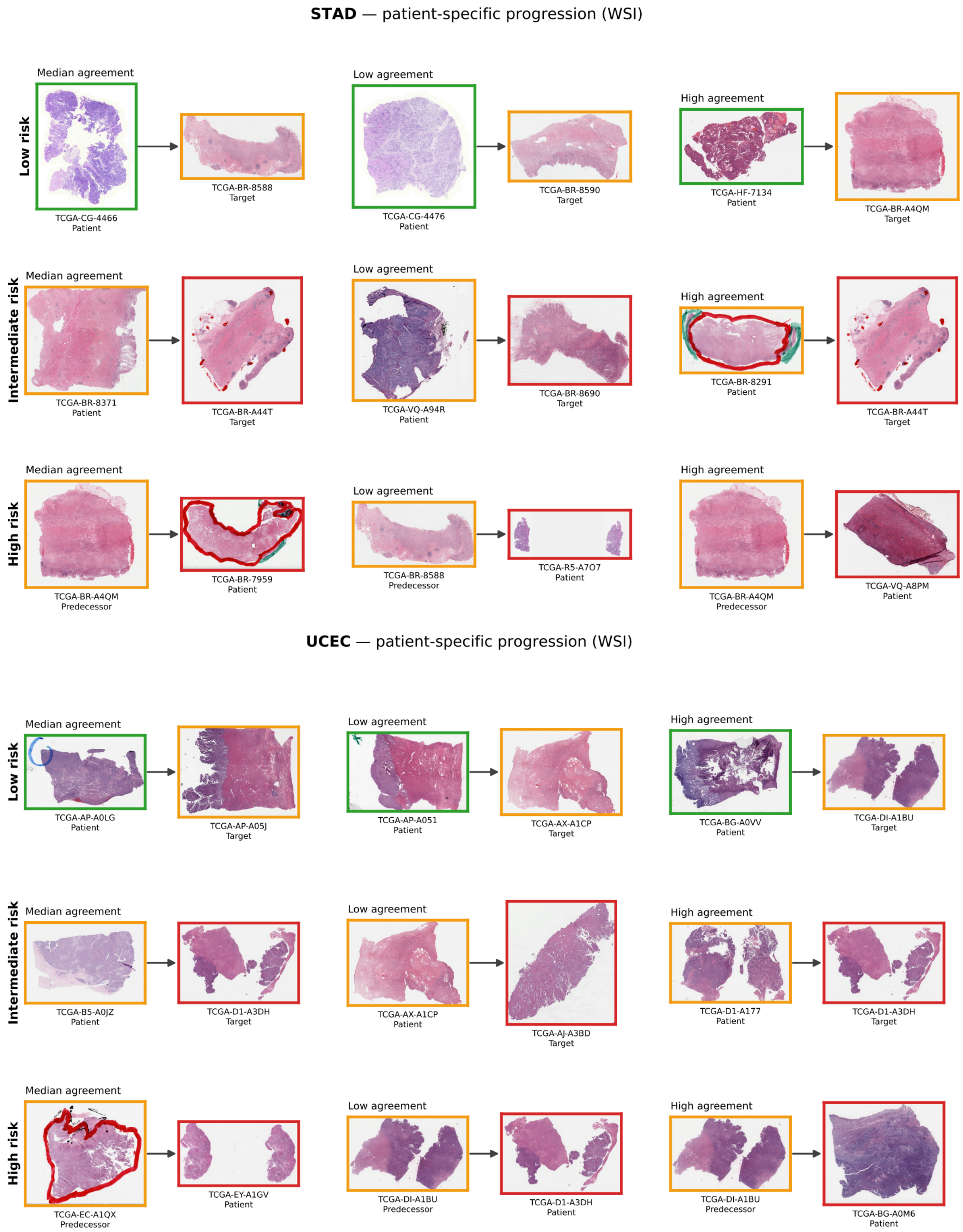


**Fig.S11.** Patient-specific progression in BLCA and CESC from whole slide images. Selected patients from each risk group are paired with their transport-derived counterpart in the neighbouring risk group. Arrows point towards higher risk, so for high-risk patients the counterpart is the inferred predecessor. Border colours denote risk group.

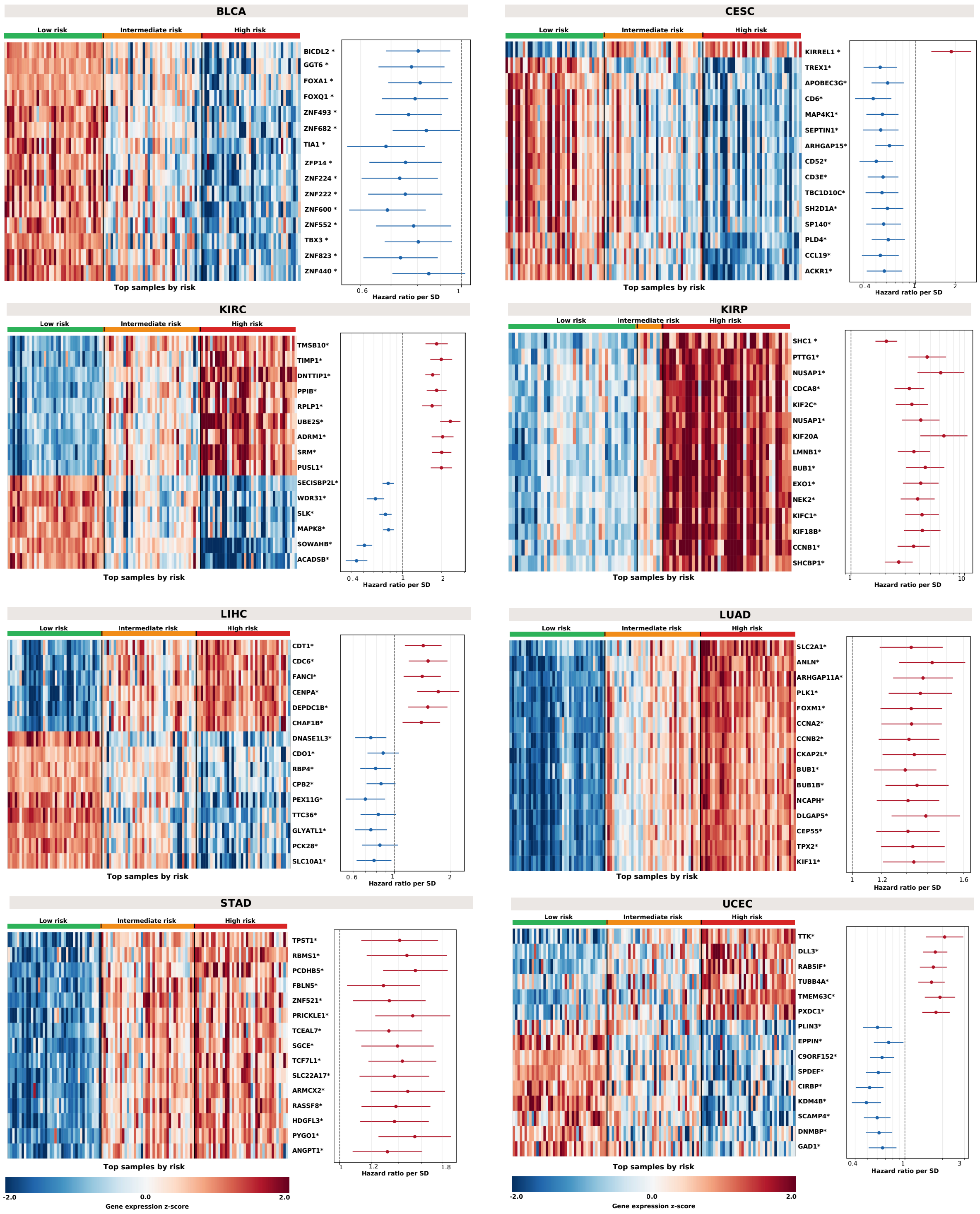


**Fig. S12.** Molecular characterisation of the GOTSurv risk groups. Heatmaps show the expression (z-score) of the genes most differentially expressed across the three risk groups (Kruskal–Wallis, Benjamini–Hochberg FDR; asterisks mark q < 0.001), with the 40 highest-risk samples per group ordered from low to high risk. Forest plots show the corresponding univariate Cox hazard ratios per standard deviation increase in gene expression, with 95% confidence intervals.

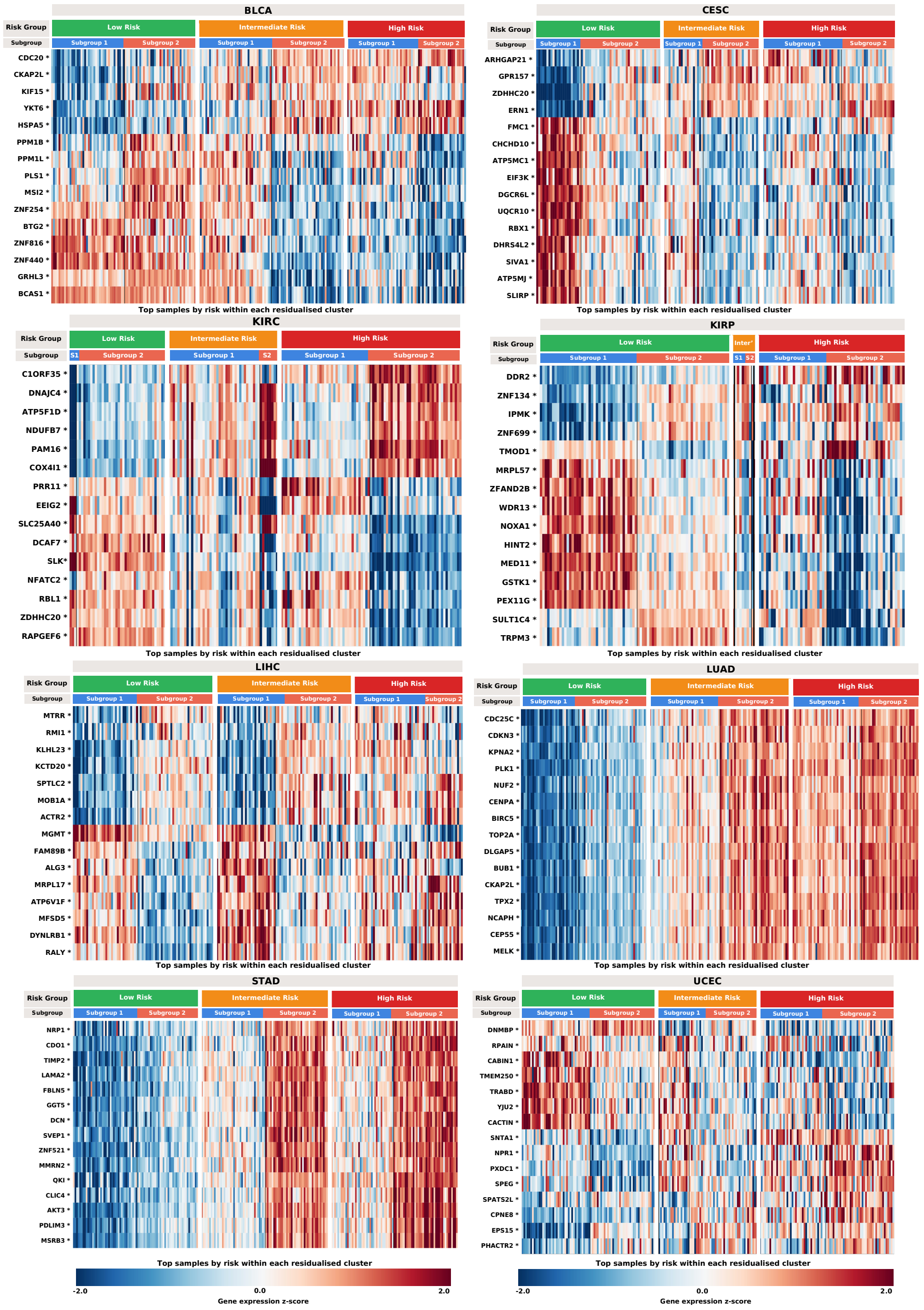


**Fig. S13.** Within-group heterogeneity of the GOTSurv risk groups. Within each risk group, samples are split into two clusters obtained from risk-residualised progression vectors, and heatmaps show the expression (z-score) of the genes best separating them (Mann-Whitney, Benjamini-Hochberg FDR; asterisks mark $q < 0.001$). Cluster blocks are ordered by expression for display only and the clustering is independent of predicted risk.

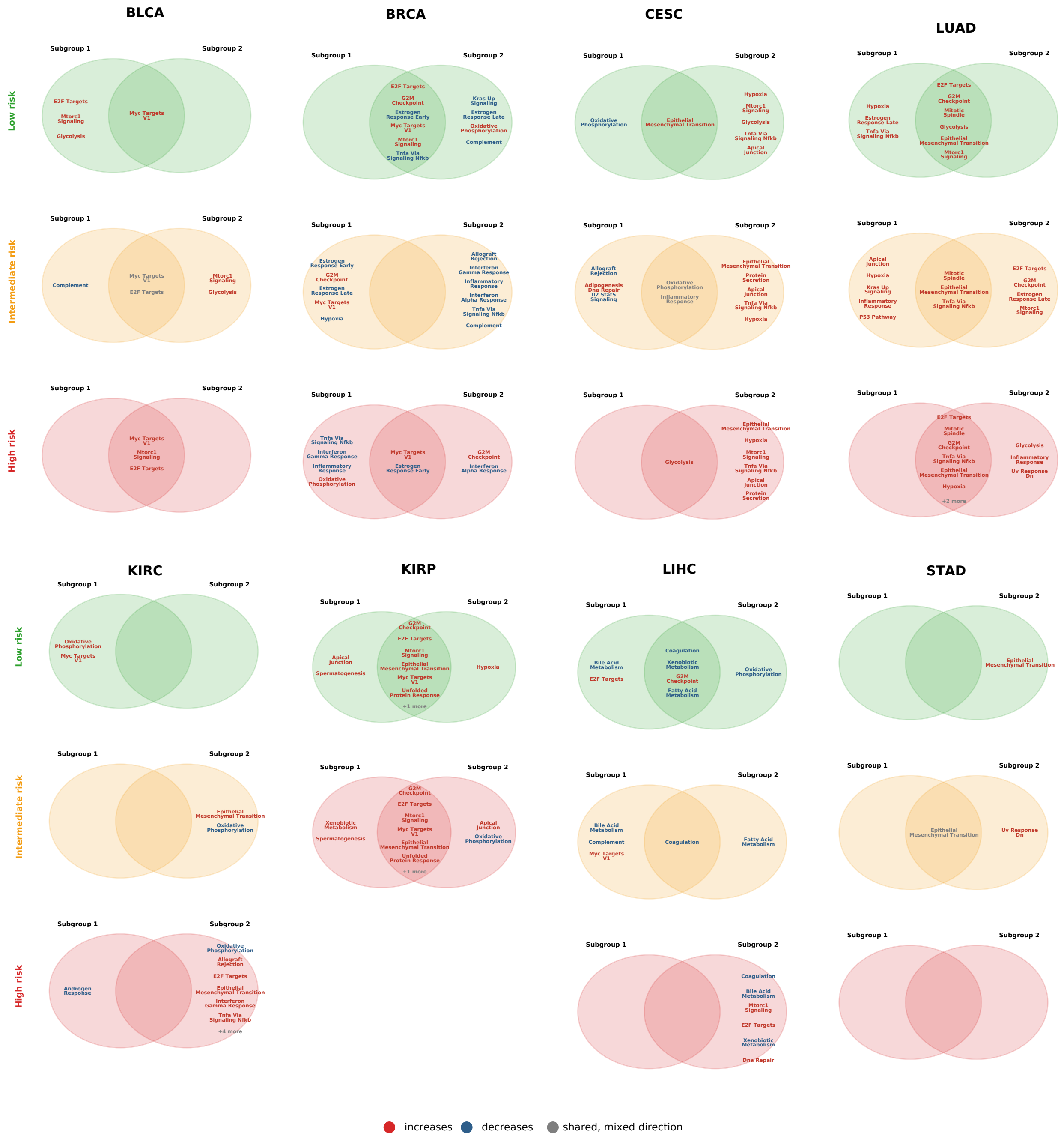


**Fig. S14.** Venn diagram showing shared and subgroup-specific progression-associated gene sets within risk groups. Shared regions indicate gene sets enriched in both progression subgroups, while non-overlapping regions indicate subgroup-specific biological programmes.

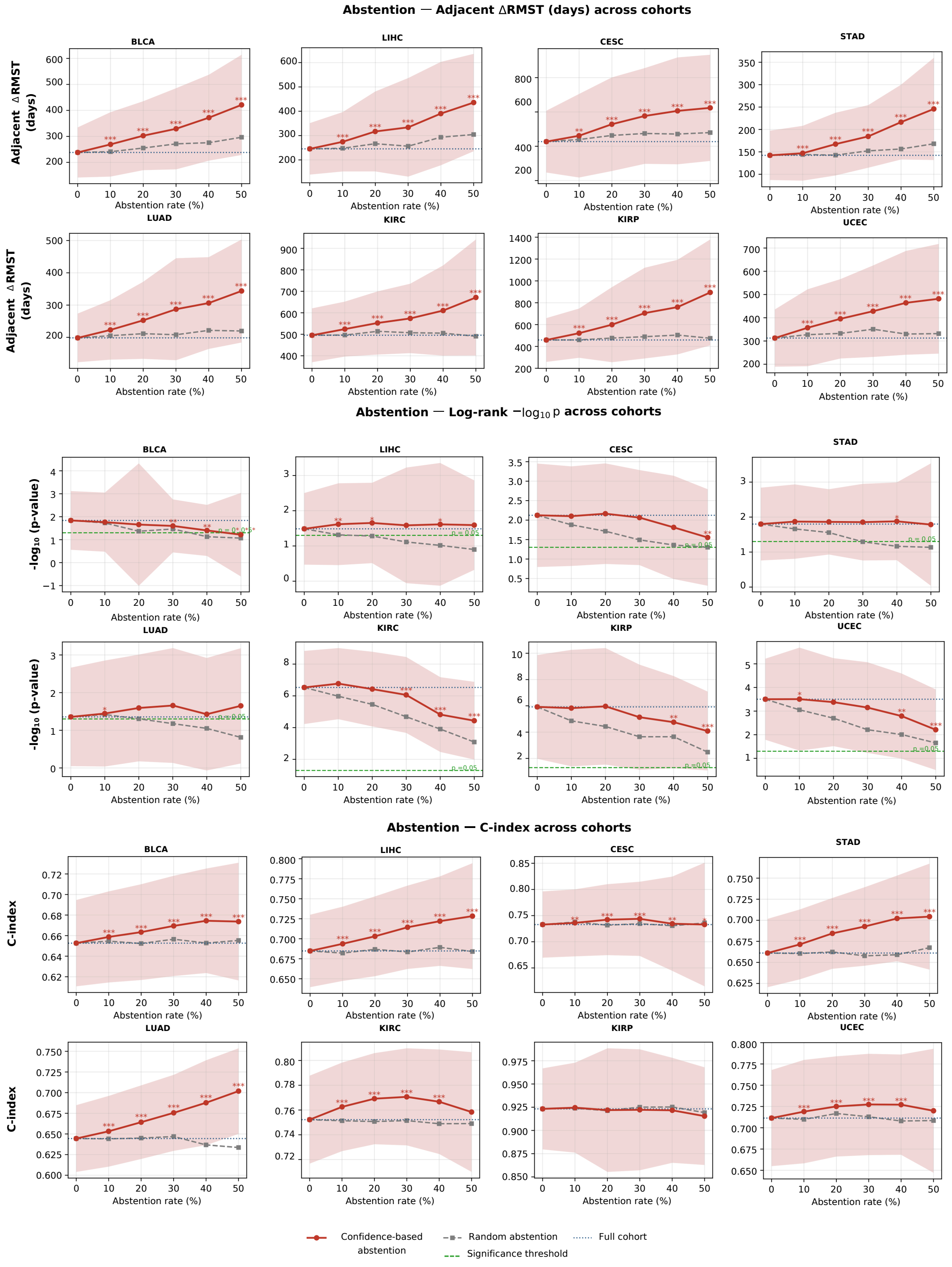


**Fig.S15.** Confidence-based abstention analysis. Changes in adjacent-group RMST separation, multivariate log-rank significance, and C-index as the rejection rate increases. Confidence-based abstention is shown against random abstention and full-cohort performance. Shaded regions show variability across resamples, and asterisks indicate significant differences from random abstention. Asterisks indicate a significant difference from the full-cohort (0% abstention) baseline using the Wilcoxon signed-rank test across matched bootstrap resamples.
